# DNA-PKcs regulates myogenesis in an AKT-dependent manner independent of induced DNA damage

**DOI:** 10.1101/2022.06.23.497315

**Authors:** Haser Hasan Sutcu, Miria Ricchetti

## Abstract

Skeletal muscle regeneration relies on muscle stem (satellite) cells. We previously demonstrated that satellite cells efficiently and accurately repair radiation-induced DNA double-strand breaks (DSBs) *via* the DNA-dependent kinase DNA-PKcs. We show here that DNA-PKcs affects myogenesis independently of its role in DSB repair. Consequently, this process does not require the accumulation of DSBs and it is also independent of caspase-induced DNA damage. We report that in myogenic cells DNA-PKcs is essential for the expression of the differentiation factor Myogenin in an Akt2-dependent manner. DNA-PKcs interacts with the p300-containing complex that activates *Myogenin* transcription. We show also that SCID mice that are deficient in DNA-PKcs, and are used for transplantation and muscle regeneration studies, display altered myofiber composition and delayed myogenesis upon injury. These defects are exacerbated after repeated injury/regeneration events resulting in reduced muscle size. We thus identify a novel regulator of myogenic differentiation, and define a differentiation phase that does not involve the DNA damage/repair process.

## Introduction

Double-strand breaks (DSBs) are dangerous DNA damages that can generate pathogenic mutations and genome rearrangements if incorrectly repaired, and compromise cell survival and cell fate if left unrepaired. The consequences of unrepaired DSBs are particularly critical in adult stem cells, which are responsible for tissue homeostasis and tissue regeneration throughout life. Unrepaired DNA damage may block proliferation, promote premature differentiation, or lead to cell death, thereby resulting in a reduction of the stem cell pool and decreased regenerative capacity of the tissue [1]. Muscle stem cells (satellite cells, SCs) are responsible for muscle regeneration and muscle homeostasis in the adult. Upon muscle damage, quiescent SCs are activated and give rise to myoblasts that differentiate and fuse into multinucleated myotubes and myofibers [2]. We previously showed that SCs repair DSBs more efficiently than their committed and differentiated progeny and that DSB repair by SCs is highly accurate, suggesting that the maintenance of genome stability is relevant for these cells [3].

The maintenance of genome stability is also important during myogenic differentiation, which is inhibited by induced genotoxic stress that blocks the activity of the myogenic determination factor MyoD [4]. Conversely, DNA damage induced by a specific DNase has been reported to promote myogenic differentiation [5]. Therefore, the role of DNA damage at different stages of myogenic differentiation needs to be clarified.

We previously demonstrated that efficient non-homologous end-joining (NHEJ) repair in SCs relies on the key repair enzyme DNA-PKcs (DNA-dependent protein kinase catalytic subunit) [3]. DNA-PKcs is recruited to DNA ends of DSBs in the presence of the Ku70/Ku80 heterodimer, then this protein kinase auto-transphosphorylates and becomes active by forming the DNA-PKcs/Ku70/80 complex called DNAPK [6]. Activated DNAPK plays a crucial role in tethering broken DNA ends [7], and acts as a signaling/mediator molecule by phosphorylating several proteins including the histone H2AX and Akt kinases thereby mediating the recruitement of these factors to the repair site, thus regulating the cellular response to the DNA lesion [8]. In addition to its direct involvement in NHEJ, upon DNA damage DNA-PKcs activates Akt kinases (Akts) prevalently located in the nucleus. Activated Akts phopsphorylate substrates that participate in DNA repair and cell cycle arrest, *i.e.* the cellular response to DNA damage.

The Akt kinases Akt1 and Akt2 are expressed in muscle cells [9]. They impact on myogenensis upon activation by PI3Ks (phosphatidyl inositol 3-kinases), which are stimulated by growth factors and cytokines [10]. PI3K/Akt1 play a role in the activation of SCs upon injury by inducing quiescence exit [11], whereas PI3K/Akt2 stimulates the expression of myogenic factors and thereby promotes muscle differentiation [12, 13]. Conversely, PIKKs (PI3K-like protein kinases) that are involved in DNA repair like DNA-PKcs and the DNA damage sensors ATM and ATR are not known to be activated during myogeneis.

We show here that DNA-PKcs affects myogenesis independently of its role in DSB repair, *via* activation of Akts. We also show that chronic impairment of this pathway *in vivo* leads to alternative activation of Akts that results in compromised muscle regeneration.

## Results

### Differentiation of satellite cells is blocked upon inhibition of DNA-PKcs

To assess the consequences of impaired DSB repair in survival and differentiation of muscle stem cells, GFP^+^ SCs isolated by FACS from *Tg:Pax7-nGFP* mice were treated for 1h with 10 µM of the DNAPK inhibitor (DNAPKi) NU7441 then exposed to 5 Gy irradiation (IR) that induces DNA damage and DSBs. Cells were kept in culture in the presence and absence of the inhibitor, and examined at 4h to 5 days post-irradiation (dpIR) (scheme Fig. 1a). Proliferation and differentiation of SCs were assessed by the cell number, or the number of nuclei in multinucleated myofibers, and immunolabeling of myogenic markers. Upon activation, Pax7^+^ myoblasts express MyoD and progressively lose Pax7 expression as they differentiate to generate Myogenin^+^ (Myog^+^) myocytes and multinucleated myofibers [14]. Myog^+^ (differentiated) cells remain MyoD^+^ for some time, then both factors are downregulated in mature myotubes.

**Figure 1.**
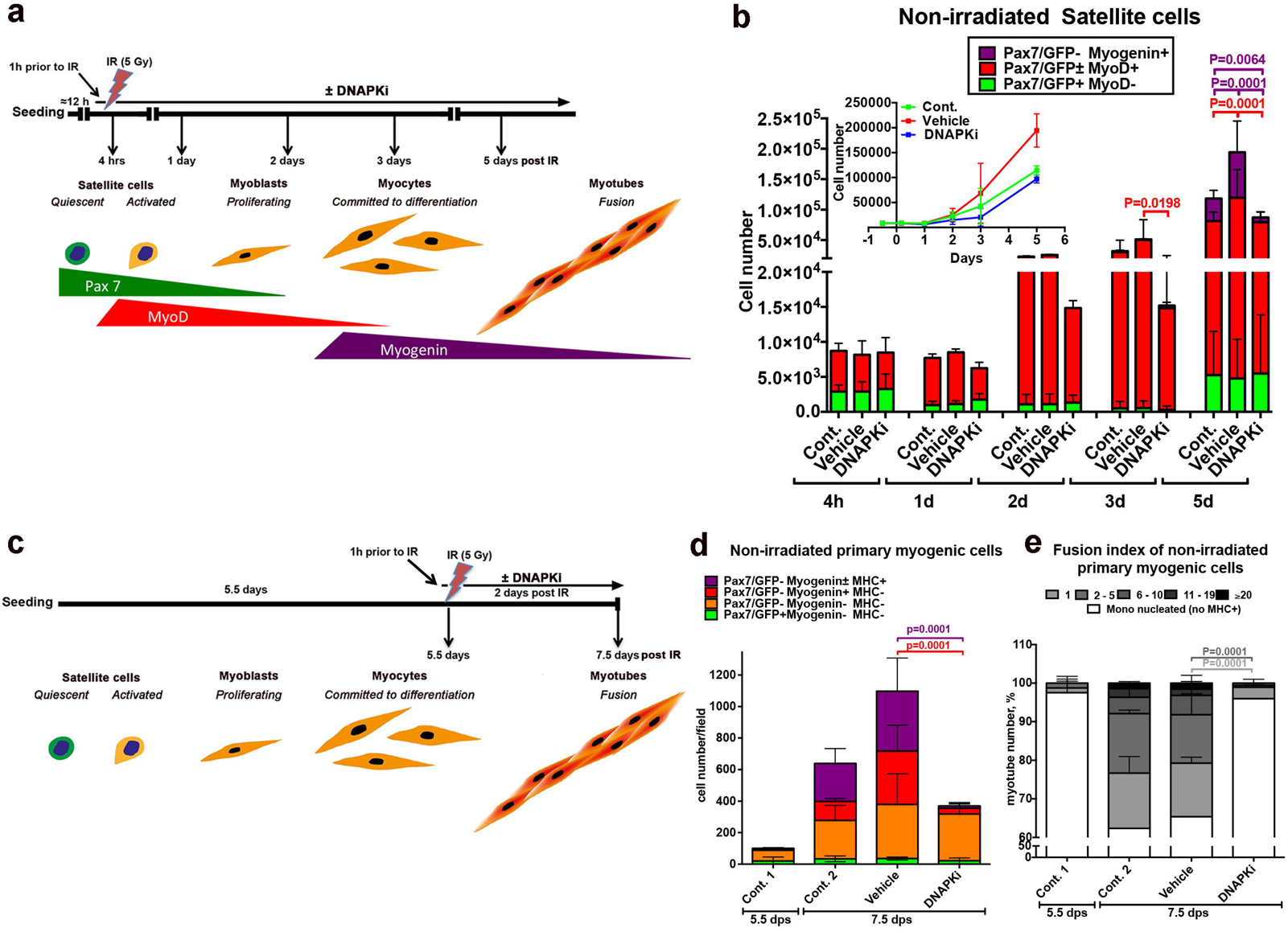
Inhibition of DNA-PKcs blocks myogenic differentiation of satellite cells in the absence of irradiation (induced DSBs) **a)** Schematic representation of the experiment: GFP^+^ SCs isolated by FACS from *Tg:Pax7-nGFP* mice were seeded overnight (≈12 hours) and pre-treated for 1h with 10 µM of DNAPKcs inhibitor (= DNAPKi= NU7441) or the corresponding volume of vehicle (0.2 % DMSO) before 5 Gy irradiation (IR), or were not irradiated. Cells were then kept in culture for 4 hours until 5 days before analysis. The expected level of differentiation and the corresponding myogenic markers at these time points are schematized below. **b)** Histograms of myogenic differentiation and growth curves of non-irradiated cells ± DNAPKi. Proliferation evaluated with the cell number, and differentiation with immunofluorescence of myogenic markers (Pax7, MyoD, Myogenin). Each condition was tested with SC derived from n=3-7 mice, mean ± SD. Myogenic markers have been analysed in 5-10 fields/condition (2,000-5,000 cells), and extrapolated to the total cell number. Significance evaluated by by 2-way ANOVA (F=7.37, DFn=28, DFd=162, p<0.000001), with post-hoc Dunnett’s multiple comparisons test; significant P values are indicated in histogram. P values and bars and are of the same colour as the category that is compared. **c)** Schematic representation of the experiment: GFP^+^ SCs isolated by FACS from *Tg:Pax7-nGFP* mice were seeded overnight and let exponentially proliferate. At 5.5 dps (days post-seeding) cells were pre-treated for 1h with 10 µM of DNAPKi or the corresponding volume of vehicle, and analysed 48h later, *i.e.* 7.5 dps. **d)** Histograms display the cell number and differentiation stage of non-irradiated SC-derived cells treated or not with DNAPKi, and defined by the indicated combinations of myogenic markers. Differentiation of MyoD^+^ myoblasts is characterized by the expression of Myogenin and later MHC. The fusion index, *i.e.* the number of nuclei per myotube, is a further parameter of differentiation at this stage. The colour code indicates the differentiation state (from Pax7/GFP^+^ [not differentiated] to MHC^+^ [the most differentiated]. At 5.5 dps, before treatments, the SC-derived population was heterogeneous (*e.g.* up to 10% [differentating] Myogenin^+^ cells, in *red* in the histogram) in agreement with published data [66]. Myogenic markers have been analysed in 5-10 fields/condition and extrapolated to the total cell number. mean ± SD for each category. Significance by 2-way ANOVA (F=2.99, DFn=9, DFd=32, p=0.0106), with post-hoc Dunnett’s multiple comparisons test; significant P values are indicated in histogram. P values and error bars are of the same colour as the category that is compared. **e)** Fusion indexes of non-irradiated SC-derived cells by enumeration of nuclei/myotube. Significance by 2-way ANOVA (F=22.68, DFn=15, DFd=35, p<0.0001.), with post-hoc Dunnett’s multiple comparisons test; significant P values are indicated in histogram. P values and error bars are of the same colour as the category that is compared. Significance bars are of the same colour as the category that is compared. n=3 experiments, n=2,500-5,000 cells analysed/condition, mean ± SD.

Irradiation decreased the number of SCs (Pax7^+^), committed (MyoD^+^) and differentiated (Myog^+^) cells, compared to non-irradiated cells (Fig. 1b and Fig. S1a, b), as expected [15]. This process was enhanced in the presence of DNAPKi, and in a dose-dependent manner (Fig. S1c, d). By 5dpIR, irradiation upon DNAPKi treatment resulted in no detectable Myog^+^ cells, and fully depleted reserve SCs (Pax7^+^/MyoD^-^/Myog^-^), that are an in vitro readout for self-renewal [16] (Fig. S1b). Impaired DSB repair upon DNAPKi treament was verified by immunostaining and enumeration of γ-H2AX foci, a histone modification prevalently associated with DSBs, and 53BP1, a protein recruited at DSBs, (Fig. S1e-g).

Surprisingly, DNAPKi also affected proliferation and differentiation of non-irradiated SCs (50% reduction of the cell number at 5 days, and 87% reduction of the Myog^+^ population compared to vehicle, Fig. 1b, and Fig. S1a), with no apparent alteration of MyoD^+^ or reserve (Pax7^+^) cells. These perturbations were observed also with the immortalized myogenic cell line C2C7, which are cycling myoblasts with high proliferative capacity (Fig. S2). These results raise the possibility that DNA-PKcs blocks myogenic cells from expressing the differentiation marker Myogenin, an activity that is independent from DSB repair.

To address the possibility that the effect of DNAPKi on myogenic differentiation is a consequence of low cell number that results from slow proliferation and/or cell death upon treatment, GFP^+^ SCs were isolated by FACS and cultured in the absence of treatment until they reached confluence (>80 %) and were poised to differentiate (Fig. 1c). At 5.5 days post-seeding (dps), cells were treated with DNAPKi (or vehicle) for 1h, then irradiated or not, and analysed two days later (*i.e*. 7.5 dps). In the absence of irradiation, treatment with DNAPKi reduced proliferation by about ≈1/3 compared to non-treated control and vehicle, where the number of Myog^+^ cells increased by 6- to 10-fold, respectively (Fig. 1d). Importantly, reduction in the cell number affected predominantly differentiated cells (Myog^+^, either MHC^-^ or MHC^+^, a terminal differentiation marker), whereas Myogenin^-^ cells were as numerous as in vehicle. Moreover, the fusion index dramatically reduced compared to vehicle (80% of the few MHC^+^ cells were mononucleated in the presence of DNAPKi whereas 63% of MHC^+^ cells contained 2-20 nuclei in the absence of treatment, Fig. 1e). These experiments show that reduced differentiation of non-irradiated myogenic cells by DNAPKi is not due to reduced cell number. As expected, irradiation affected differentiation to a larger extent, in particular when DNA repair was inhibited, upon DNAPKi treatment (Fig. S3a-c).

These results were confirmed with C2C7 myoblasts using two distinct protocols that mimic either the conditions of highly proliferative SCs (Fig. S3d-g) or a later stage with myogenic cells ready to differentiate (Fig. S3h-k), before treatment (± DNAPKi and ± irradiation). In both cases, DNAPKi treatment inhibited Myog expression in non-irradiated cells. These data also reveal that DNAPKi affects mostly differentiating (Myog^+^) rather than committed (MyoD^+^) cells, and blocks further steps of differentiation in the absence of induced DNA damage.

### Inhibition of DNA-PKcs blocks Myogenin expression in Akt-dependent manner

To verify that DNAPKi affects Myogenin expression, we treated C2C7 cells with increasing doses of DNAPKi (5-20 µM), following the previous scheme (Fig. S3d). We also examined the effect on Myogenin expression of inhibitors of PI3Ks (PI3Ki; ZSTK474, or LY294002), since DNA-PKcs is a PI3K-related kinase [17] [18], as well as an inhibitor of the downstream Akt kinases (AKTi; MK2206). Akt kinases are activated (phosphorylated at Ser473 and Thr308) by PI3Ks, as well as by PIKKs, upon DNA damage[19].

The transcript levels of *Myog* were reduced by 88% in cells treated with 10 µM DNAPKi compared to vehicle at 5 dps (Fig. 2a) and, accordingly, the Myogenin protein signal was depleted in a dose-dependent manner (Fig. 2b). The block in Myogenin expression was associated with reduced phosphorylation of Akts (phospho-Akts, or p-Akts) (Fig. 2c), indicating that DNAPKi also acts on Akts activation. Further, treatment with PI3Ki to some extent reduced, and AKTi depleted, Myogenin expression (Fig. 2e). Importantly, MyoD was not reduced with either of these three inhibitors (Fig. 2b). Depletion of Myogenin, reduction of p-Akts with increasing doses of DNAPKi, as well as reduced levels of Akts, were all confirmed in SCs (Fig. 2d).

**Figure 2.**
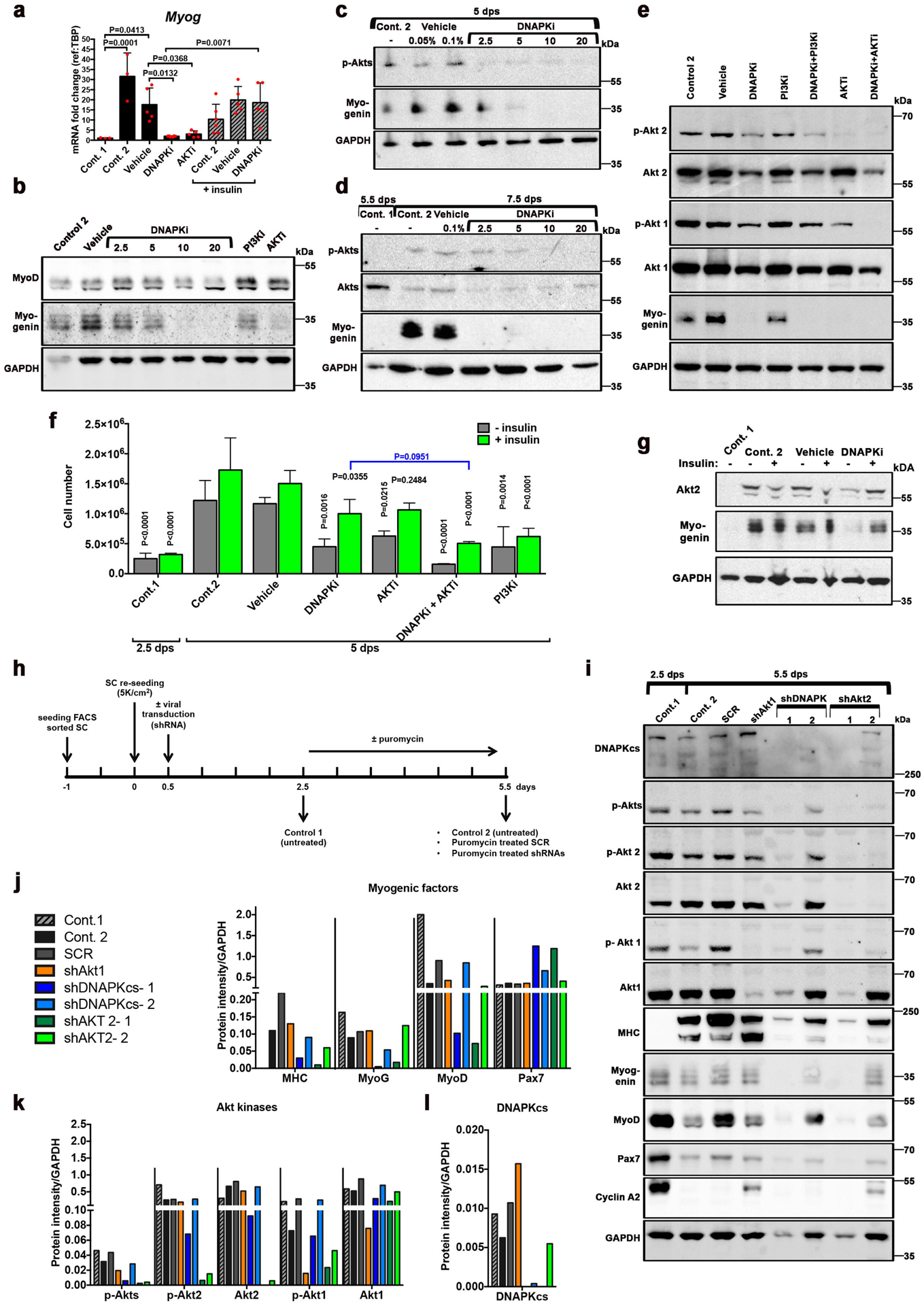
DNA-PKcs is required for expression of myogenin in a Akt-2 dependent manner. **a-g)** Myogenic cells treated with inhibitor of kinases. C2C7 cells were grown until at 2.5 dps, following the scheme in Fig. S3d, and treated with inhibitors (alone or in combination, at the indicated doses) until 5 dps (panels a-c, e). SCs (panel d) were treated following the scheme in Fig. 1c, *i.e.* treated at 5.5 dps and collected at 7.5dps. Control 1, untreated at 2.5 dps (or 5.5 dps, panel d); control 2, untreated at 5 dps (or 7.5 dps, panel d); vehicle, untreated (DMSO) at 5 dps (or 7.5 dps, panel d). **a)** mRNA fold changes of *Myog* expression (n=3-6) in C2C7 cells, mean ± SD normalised to TBP-housekeeping gene, black columns, left part of the histogram. In the righ part of the histogram, grey hatched columns indicate the same samples in the presence of (10 µg of insulin). Significance by Ordinary 1-way ANOVA (F=9.943, DFn=7, DFd=30, p<0.0001) with post-hoc Tukey’s multiple comparisons test, significant p-values are indicated on the histogram. **b)** Western blot (WB) of C2C7 cells treated with increasing doses of DNAPKi (NU7441), or 1 µM PI3Ki (ZSTK474), or 5 µM AKTi (MK2206); two bands of MyoD are present [67], and a band shift is observed at the highest doses of DNAPKi. **c**) WB of Myogenin and p-Akts in C2C7 cells at increasing concentrations of DNAPKi. Two vehicle lanes (0.05% and 0.1%) correspond to the volume of DMSO used with 2.5 µM and 10 µM DNAPKi, respectively. **d)** WB of Myogenin, Akts, and p-Akts of SCs in culture until 5.5 dps before treatment with increasing concentrations of DNAPKi for two more days. **e)** WB of C2C7 cells treated with 10 µM DNAPKi, PI3Ki (LY294002), and 5µM AKTi, alone or in combination (left panels), and cultured in the presence or in the absence of inhibitor(s) until 5 dps; in panels b and e, vehicle contains 0.1% DMSO. For WBs, GAPDH was used as reference housekeeeping protein. **f)** C2C7 cell number and **g)** WB of Myogenin and Akt2, upon growth and differentiation in the presence and in the absence of various inhibitors and insulin (green columns). As expected, insulin did not improve proliferation of cells treated with PI3Ki, because PI3Ks are inhibited. In WB GAPDH was used as reference housekeeeping protein. Significance by 2-way ANOVA (F=4.66, DFn=18, DFd=69, p<0.0001), with post-hoc Tukey’s multiple comparisons test vs. corresponding vehicle conditions (vehicle or vehicle + insulin) when not specifically indicated (P values shown on the histogram). **h-l)** Analysis of Myogenic factors and Akt kinases upon shRNA-dependent silencing of Akt1, Akt2 or DNA-PKcs. **h**) Scheme and readouts of the experiment: SCs tested at 2.5 dps and 5.5 dps (control 1 and control 2, respectively) in the absence of puromycine (selection) treatment; SCR (scramble RNA) or shAkt1, shDNA-PKcs (2 independent clones), and shAkt (2 independent clones) at 5.5 dps upon puromycine selection. **i**) WB of myogenic factors (differentiation markers: MHC (MF20) and Myogenin; myoblast marker MyoD, and stem cell marker Pax7), kinases (Akt1, Akt2, phospho-Akt1, phosphor-Akt2, phospho-Akts, and DNA-PKcs), and the cell cycle marker Cyclin A2. MyoD levels remained relatively high in SCR, perhaps as a consequence of the lentiviral and selection procedure. Normalisation of protein levels with the reference protein GAPDH upon quantification of bands with Imagelab (BIORAD) for **j**) myogenic factors, **k**) individual Akt kinases and their phosphorylated forms, and global p-Akts, and **l)** DNA-PKcs. Uncropped gels are shown in Fig S5.

Both members of the Akt family that are expressed in muscle (Akt1 and Akt2 [9]), and their respective phopsphorylated forms, and in particular p-Akt2, were affected by DNA-PKcs inhibition (Fig. 2e). Conversely, PI3Ki, which moderately reduced Myogenin levels, poorly reduced p-Akt2, and did not affect the levels of Akt1, p-Akt1, and Akt2, confirming a limited effect of PI3Ks on myogenesis. Double treatment with DNAPKi and PI3Ki resulted in a larger reduction of p-Akt2 than DNAPKi alone suggesting that activation of Akt2 by DNA-PKcs and PI3Ks are at least in part, non-overlapping processes. Conversely, this double treatment did not change the levels of Akt1/pAkt1 and global Akt2 compared to DNAPKi alone, confirming the above-mentioned lack of effect of PI3Ki on these factors.

Finally, AKTi that inhibits Akts phosphorylation by reducing the levels of p-Akt1 and almost suppressing p-Akt2, inhibited Myogenin expression thus confirming that this effect depends on both Akts activation and DNA-PKcs (Fig. 2e). Combined AKTi and DNAPKi treatment further reduced the levels of Akt1 and Akt2, above that of each single inhibitor alone, confirming the notion that Akt1 and Akt2 phosphorylation during early myogenesis depends to a large extent, but not exclusively, on DNA-PKcs.

In summary, experiments with multiple inhibitors of kinases indicate that DNA-PKcs affects the fate of myogenic cells by promoting Myogenin expression, and this depends on activation of both Akt1 and Akt2. Conversely, PI3Ks have a minor effect and only result on moderate Akt2 activation.

### Insulin-induced differentiation alleviates DNAPKi-blocked myogenesis through the PI3K/Akt pathway

Insulin and insulin-like growth factors (IGF) regulate and activate PI3Ks [20]. Insulin was added to the DNAPKi treatment, to assess whether insulin-dependent PI3K activation of Akts, and consequently inhibition of myogenesis, were rescued. C2C7 cells were treated with DNAPKi and/or inhibitors of the PI3K/Akt pathway (PI3Ki or AKTi), in the presence and in the absence of 10 µg/ml of insulin. Insulin (green columns, Fig. 2f) rescued to a large extent proliferation of cells treated with either DNAPKi or AKTi by doubling the cell number, and displayed its largest effect (3-fold increase) in the presence of the two combined inhibitors, *i.e.* the condition in which proliferation was the most significantly reduced. Consistent with our model, insulin partially restored the levels of Myogenin transcripts (hatched columns Fig. 2a, and protein (Fig. 2g) that were significantly reduced upon DNAPKi treatment. Moreover, insulin rescued the levels of Akt2 that was also depleted upon treatment with DNAPKi (Fig. 2g). Thus, insulin alleviates DNAPKi-dependent/AKTi-dependent depletion of Myogenin and the decline in cell proliferation, suggesting that Myogenin expression can be, at least partially, activated by PI3Ks, when DNA-PKcs is inhibited.

### Myogenin expression is blocked by silencing DNA-PKcs and Akt2

To confirm that impairment of myogenic differentiation by DNAPKi and AKTi is due to *bona fide* inhibition of DNA-PKcs and Akt, the expression of DNA-PKcs, Akt1, and Akt2 was independently silenced in SCs by shRNAs lentivirus transduction (Fig. 2h). Control 1 (cells harvested 2.5 dps, in absence of treatment) displayed strong Pax7 and MyoD, and little Myogenin signal (Fig. 2i, j), confirming the prevalent myoblast state; the proliferation of these cells was verified by high levels of cyclin A2, a marker of dividing cells. At 5.5 dps, as expected, cyclin A2 levels dropped in control 2 (untreated) and the SCR sample (scramble shRNA). Advanced differentiation of both cells was evidenced by high levels of MHC, concomitantly with reduction of Myogenin and MyoD, as well as depletion of Pax7.

Silencing DNA-PKcs (by >95%) with two independent clones resulted in strongly reduced or undetectable MHC and Myogenin (with a larger effect with shDNAPK-1) compared to SCR, confirming that DNA-PKcs is necessary for myogenic differentiation (Fig. 2i, j). Silencing with shDNAPK-1 decreased MyoD levels, indicating that the committed myogenic population was also affected by DNA-PKcs knock-down. The more efficient silencer, shDNAPK-1, reduced the levels of globals p-Akts, and specifically Akt1, Akt2 and the respective phsophorylated forms (Fig. 2i, k).

Akt1silencing (by >90% with shAkt1) had a weak effect on committed myogenesis as it essentially affected myoblasts (Fig. 2i, j), in agreement with previous findings [9], and reduced MHC but not Myogenin levels. Conversely, Akt2 silencing with either shRNAs reduced the levels of MHC and MyoD as much as silencing of DNA-PKcs; the levels of Myogenin were reduced only by shAkt2-1. Of note, silencing of Akt2 also depleted DNA-PKcs (Fig. 2i, l).

Altogether, silencing experiments confirmed that DNA-PKcs is implicated in myogenesis and showed that the DNA-PKcs downstream target Akt2 has similar effect on myogenesis as DNA-PKcs itself, whereas Akt1 only mildly affects this process.

### DNA-PKcs interacts with the complex that activates Myogenin expression

DNA-PKcs may regulate Myogenin expression by direct intervention on its promoter or activating one of the effectors of the Myogenin promoter. Chromatin immunoprecitation (ChIP) experiments on the Myogenin promoter region confirmed the expected presence of MyoD [21, 22] but not DNA-PKcs, which is instead present on the locus only upon DNA damage (positive control; Fig. 3a-c). These experiments suggest that DNA-PKcs does not directly regulate the Myogenin promoter. Conversely, co-immunprecipitation experiments with DNA-PKcs pull-down revealed the presence of MyoD and the acetyltransferase p300 (Fig. 3d, e), two key components of the complex that activates myogenic genes including Myogenin [21], suggesting that DNA-PKcs is part of the same complex as these activators of Myogenin. P300 and MyoD were also present upon Akt1 pulldown (Fig. 3d), and p300 upon Akt2 pulldown, indicating that p300 and MyoD containing complexes interact with Akts as well, consistent with Akts phosphorylating p300 to promote the p300/MyoD interaction [21]. Finally, Akt1 and Akt2 were present upon DNA-PKcs pulldown (Fig. 3f).

**Figure 3.**
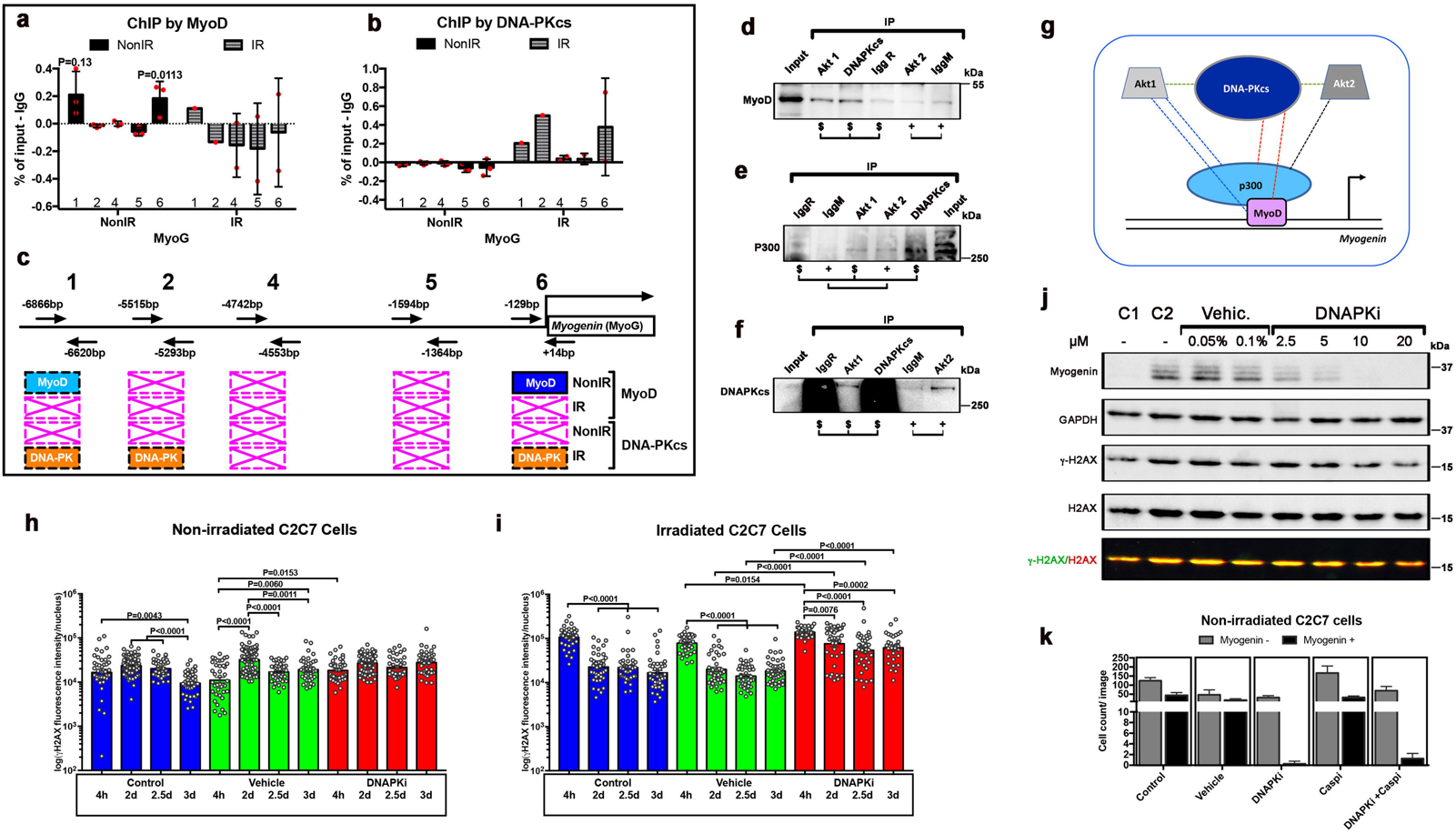
DNA-PKcs interacts with the complex that activates Myogenin expression, and activates Myogenin/myogenic differentiation without acting in DNA repair. Quantitative PCR (histogram) analyses of several DNA fragments in the *Myogenin* promoter [68] from ChIP assays with either **(a)** anti-MyoD or **(b)** anti-DNA-PKcs antibodies in differentiating cells Significance by Unpaired t-test (F=2.9, DFn=2, DFd=2). **(c)** Scheme not at scale of primer positions on the the *Myog* promoter wih the respective distances from the initiation of transcription site (+1), and summary of positive amplifications in light blue and statistically significant in dark blue (MyoD) or orange (DNA-PKcs) whereas crossed boxes in pink indicate no amplification, in the absence and in the presence of irradition: n=3 unless diffferently indicated. Immunostaining with **(d)** MyoD, **(e)** p300, and **(f)** DNA-PKcs of immunoprecipitation with DNA-PKcs and Akt1 (IgG rabbit, IgR), and Akt2 (IgG Mouse, IgM) pull-down. Input, empty IgG rabbit (IgGR, for control of DNA-PKcs and Akt1) and IgGM (IgG Mouse, IgM, for control of Akt2) in C2C7 cells at 5dps. **g)** Schematic representation of the interactions (dotted lines) observed in immuniprecipiatation experiments; dotted lines of the same colour indicate interactions detected with pulldown of the same protein. Protein sizes are not at scale. Histone γH2AX immunostaining evaluated *per* cell in the absence [control, vehicle (0.2% DMSO)] and in the presence of 10 µM DNAPKi at different time points **h)** in non-irradiated cells and **i**) irradiated C2C7 cells. Total γH2AX immunofluorescence intensity was assessed with ImageJ of single cells after acquisition with Cell Voyager CV1000, confocal scanner box (Yokogawa). N= 40-80 cells from 3 independent experiments. Significance by 1-way ANOVA (Non-irradiated: F=16.37, DFn=11, DFd=588, p<0.0001. Irradiated: F=53.68, DFn=11, DFd=457, p<0.0001), with post-hoc Tukey’s multiple comparisons test. P-values are indicated on the histogram. **j)** WB of Myogenin, global histone H2AX, γH2AX, and the housekeeping GAPDH protein in C2C7 cells at 4.5 dps. The ratio γH2AX/H2AX is shown in the bottow line after superimposition of the (700 nm (Goat α-mouse Starbright blue 700) and 800nm (goat α-rabbit CF770) signal with the imagelab program of ChemiDoc MP imaging system (Biorad). The highest doses of DNAPKi results in slightly reduced levels of γ-H2AX, compatibly with H2AX being also phosphorylated by DNA-PKcs. **k**) Enumeration of Myogenin^+^ and Myogenin^-^ cells upon immunostaining per image in the presence and in the absence of inhibitors of DNA-PKcs and caspases (4 images/condition analysed, n=1). Uncropped gels are shown in Fig S5.

Altogether, these data indicate that DNA-PKcs interacts with p300 and MyoD that mediate Myogenin expression at its promoter, as well as its regulators Akt1 and Akt2 (recapitulative scheme in Fig. 3g). Whether DNA-PKcs participates in these complexes in the context of DNA repair remains unclear.

### DNA-PKcs is required to activate Myogenin-dependent differentiation without acting on DNA repair

Since DNA-PKcs is normally activated by DNA damage, DNA-PKcs function in promoting myogenesis could be triggered by endogenous DNA damage, for instance generated during DNA replication or, alternatively, by caspase-induced DNase (CAD). CAD-generated DNA damage at promoters has been reported to induce the expression of genes such as p21, which blocks the cell cycle, to promote late muscle differentiation [5]. In the same study, direct DNA damage on the *Myogenin* locus was not observed. We reported above DNA damage in the absence of irradiation in SCs (Fig. S1e-g) and activated C2C7 myoblasts (not shown) up to 24h and 48h in culture, respectively, which is compatible with DNA lesions induced during DNA replication [23, 24]. Analysis of DNA damage at later time points showed that in the absence of irradiation, the γ-H2AX signal/cell remained essentially constant after 3 days in culture (Fig. 3h), a time point when Myogenin starts to be expressed in C2C7 cells and DNAPKi affects myogenesis (see above Fig. S2c). The γ-H2AX signal was not affected by DNAPKi (Fig. 3h). This result indicates a nearly constant load of potentially replication-induced DNA damage that is not repaired by DNA-PKcs, as expected for this type of DNA lesion [23, 24]. As a control, DNAPKi largely impaired the repair of IR-induced DNA damage in the same culture conditions (Fig. 3i). Consistent with this result, WB analysis confirmed that γ-H2AX is present in C2C7 myoblasts at 2.5 dps (control 1) as well as during differentiation (control 2), and did not increase in the presence of DNAPKi (Fig 3j). These data suggest that DNA-PKcs is not activated for the repair of endogenous DNA damage during differentiation. Also, DNA-PKcs is not involved in phosphorylation of H2AX in response to replication-induced DNA damage, which substantially does not vary during differentiation.

DNA-PKcs may then be involved in myogenesis through processing DNA damage produced by other sources, *e. g.* CAD, where DNA-PKcs has been previously reported to phosphorylate H2AX [25]. However, differently from that study, inhibition of caspases by CASPi did not reduce the number of Myogenin expressing cells, whereas DNAPKi fully blocked the appearance of these cells (Fig. 3k). These data suggest that in our model DNA-PKcs impairment affects muscle differentiation independent of, and possibly acts at an earlier stage of myogenesis than caspases.

### Satellite cells from SCID mice have reduced proliferation and differentiation

To evaluate whether altered myogenesis affects muscle regeneration in the context of chronic DNA-PKcs depletion, we analysed SCs derived from SCID mice that are naturally DNAPK-deficient, and control mice on the same genetic background (BALB/c) [26]. Freshly isolated SCs that were plated and assessed from 1.5 to 7.5 dps displayed comparable proliferation rates in WT and SCID mice until 5.5 dps, although at this time point about 10 % less Myog^+^ cells were present in SCID-derived cells (Fig. 4a, b). Moreover, the fusion index (MHC staining) showed about 80% mononucleated MHC^+^ cells in SCID *versus* 60% of mononucleated and 40% multinucleated cells in WT (Fig. 4c). No delay in myofusion in mutant cells was observed at 7.5 dps (Fig. 4c), but we detected 30 % less reserve SCs in SCID compared to WT (light green, Fig. 4b), indicating that not only differentiation but also self-renewal of SCs was perturbed in SCID-derived cells. Thus, chronic DNA-PKcs deficiency in mice affects myogenesis, but to a lower extent than the DNAPKi treatment or DNA-PKcs silencing, suggesting a compensatory mechanism as these mice develop muscles in the absence of DNA-PKcs.

**Figure 4.**
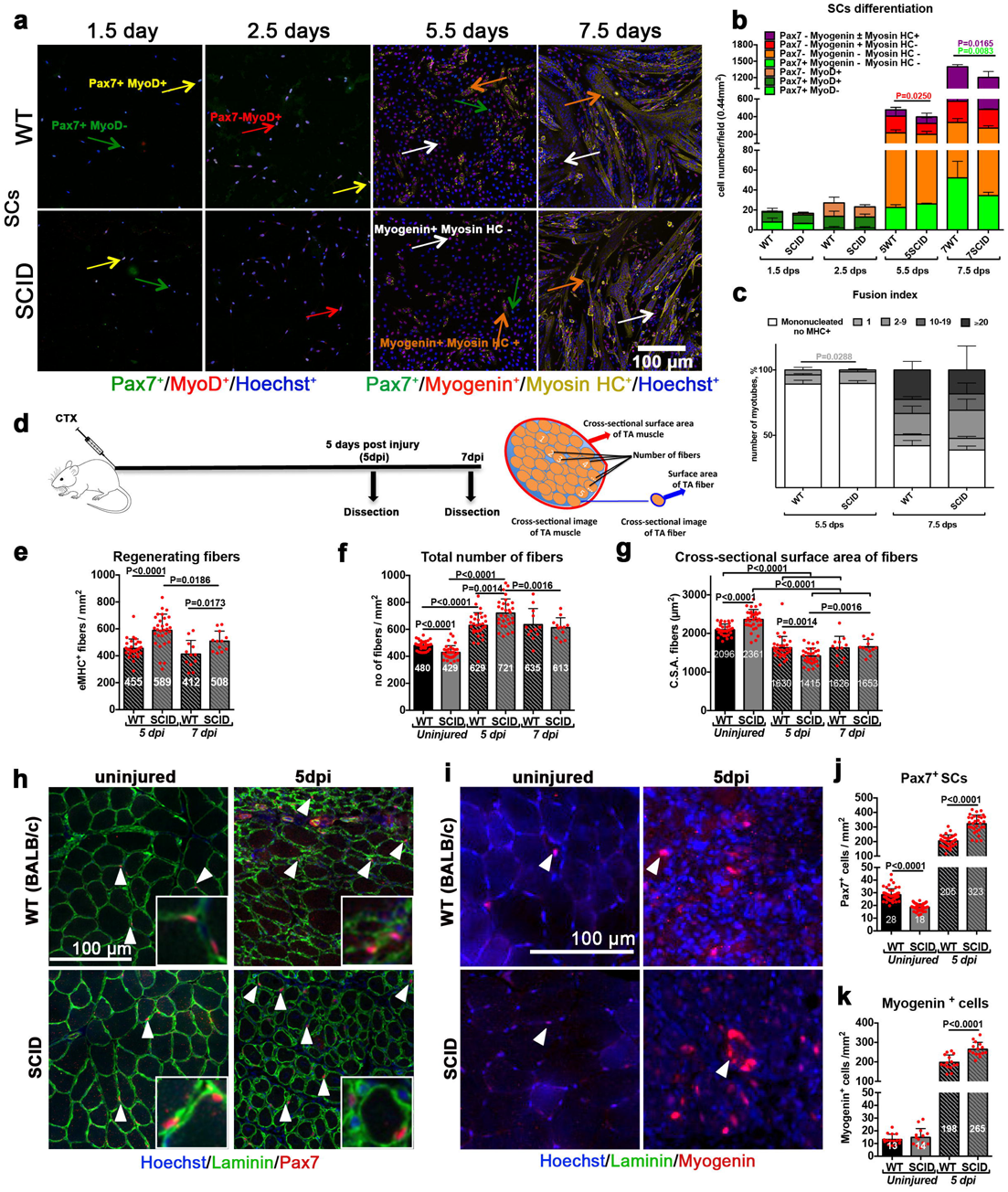
Delayed SCs differentiation in SCID mice results in altered muscle composition during regeneration. (a-c) SCs isolated from SCID and the corresponding WT mice. **a)** Immunofluorescent labelling of WT or SCID SC-derived myogenic cells with: Pax7 (green), MyoD (red), Myogenin (red), and MHC (yellow); nuclei are counterstained with Hoechst (blue). Pax7^+^ cells are indicated with a green arrow, MyoD^+^ cells with a red arrow, Myogenin^+^ cells with a white arrow, and MHC^+^ cells with an orange arrow. **b)** Histograms of myogenic differentiation of SC derived from WT and SCID mice. Combination of myogenic markers reveals several myogenic or differentiated subpopulations, indicated in the legend of the histogram, and displayed with increasing differentiation level from bottom to top. n=3-4 mice, 5-10 fields/condition, mean ± SD. Significance by Mann Whitney test, P values indicated on the histogram. **c)** Fusion index 5.5 and 7.5 days post-seeding. The number of nuclei/structures have been arbitrarily defined and vary from 0 to >20. Sixty-seven then 789 MHC^+^ cells/0.44 mm^2^ in SCID mice and 73 then 870 MHC^+^ cells/0.44 mm^2^ in WT mice at 5.5 dps and 7.5 dps respectively. n=3-4 mice, 5,000 nuclei/condition, mean ± SD. Significance by Mann Whitney test, P values are indicated on the histogram. (d-h) *In vivo* TA muscle regeneration in WT and SCID mice 5 and 7 dpi. **d)** Experimental scheme: 10 µM of cardiotoxin injury followed by TA dissection 5 and 7 dpi, along with un-injured TA from the same mouse. Cross sectional scheme of TA muscle and parameters measured are shown. Histograms of **e)** number of regenerating eMHC^+^ fibers/mm^2^, **f**) number of total fibers/mm^2^, and **g)** cross-sectional surface area (C.S.A.) of fibers on TA sections. The C.S.A. of the fibers is a measure of their size (thickness), obtained by dividing the TA surface by the number of fibers. Representative images of e, f and g are shown in Fig. S4a. Representative images of **h)** Pax7^+^ SCs (in red) and **i)** Myogenin^+^ differentiating cells (in red), indicated with arrowheads on un-injured and 5dpi TA sections. Quantification of **j)** Pax7^+^ SCs/mm^2^ and **k)** Myogenin^+^ cells/mm^2^ on un-injured and 5dpi TA sections. n=3 mice, 10 sections/condition, mean ± SD. Significant P values between conditions by Mann Whitney test are indicated on the histograms.

### Altered myofiber composition and structure after regeneration in SCID mice

To assess regeneration of DNA-PKcs deficient SCs *in vivo*, SCID and the corresponding WT control mice were injured with cardiotoxin (CTX) in the *Tibialis anterior* (TA) muscle. TA muscles were dissected and analyzed 5 and 7 days post injury (dpi) (Fig. 4d). Regeneration was assessed by quantification of embryonic eMyHC^+^ fibers (Fig. S4a), a marker that is expressed transiently during muscle regeneration [27]. We observed a higher density of eMyHC^+^ fibers in injured SCID compared to WT TA muscle (589 and 459 eMHC^+^ fibers/mm^2^ at 5dpi respectively, Fig. 4e), and consequently a higher density of total fibers (enumerated after Laminin^+^ labelling; 721/mm^2^ in SCID and 628/mm^2^ in WT; Fig. 4f). Although the number of eMyHC^+^ regenerating fibers decreased from 5dpi to 7dpi, they remained at higher density in injured SCID than WT TA muscle (508 and 412 eMyHC^+^ fibers/mm^2^, respectively at 7dpi; Fig. 4e), and the total number of fibers was not significantly different at 7dpi (613 fibers/mm^2^ in SCID TA and 635 fibers/mm^2^ in WT TA; Fig. 4f). In contrast, in uninjured TA muscle the density of total fibers was higher in WT than SCID mice (480/mm^2^ and 429/mm^2^, respectively, Fig. 4f). To uncouple the regeneration defects due to DNA-PKcs deficiency from immunodeficiency [28], we also injured *Rag2^-/-^γ c^-/-^* immunodeficient mice and the corresponding control (C57BL/6), and observed no difference in the density of regenerating fibers at 5dpi (Fig. S4b).

SCID fibers displayed a smaller (16%) cross-sectional area (CSA) than WT at 5 dpi, and this difference was lost at 7dpi (Fig. 4g). In contrast, in uninjured mice the CSA of fibers was 10% larger in SCID than WT mice. In summary, SCID mice have slightly less fibers, but these fibers cover a larger surface than in WT. Upon injury, the situation is reversed and fibers increase in number and cover a smaller surface in SCID than in WT mice during early regeneration, then recover later, pointing to a delay in muscle regeneration in SCID mice.

Cross-sectional area of the entire TA muscle showed that that uninjured SCID have less Pax7^+^ SCs than WT (18 cells/mm^2^ and 28 cells/mm^2^, respectively) (Fig. 4h, j). This situation was reversed after CTX injury (323 Pax7^+^ SCs cells/mm^2^ in SCID *vs.* 205 Pax7^+^ SCs cells/mm^2^ in WT TA). Similarly, differentiating (Myogenin^+^) cells were more numerous in injured SCID (265/mm^2^) than WT TA muscle (198/mm^2^) (Fig. 4i, k). Therefore, SCID mice show hallmarks of delayed regeneration that are compensated at 7 dpi.

### Repetitive injuries accentuate regeneration defects in SCID mice

We then asked if SCID mice can sustain multiple cycles of regeneration, as is the case for WT mice [26, 29]. To do so, TA muscles were injured with CTX from 1 to 3 consecutive times, allowing 30 days for regeneration for each cycle [26]. Three sets of mice were used for one, two, and three rounds of injury, respectively (Fig. 5a).

**Figure 5.**
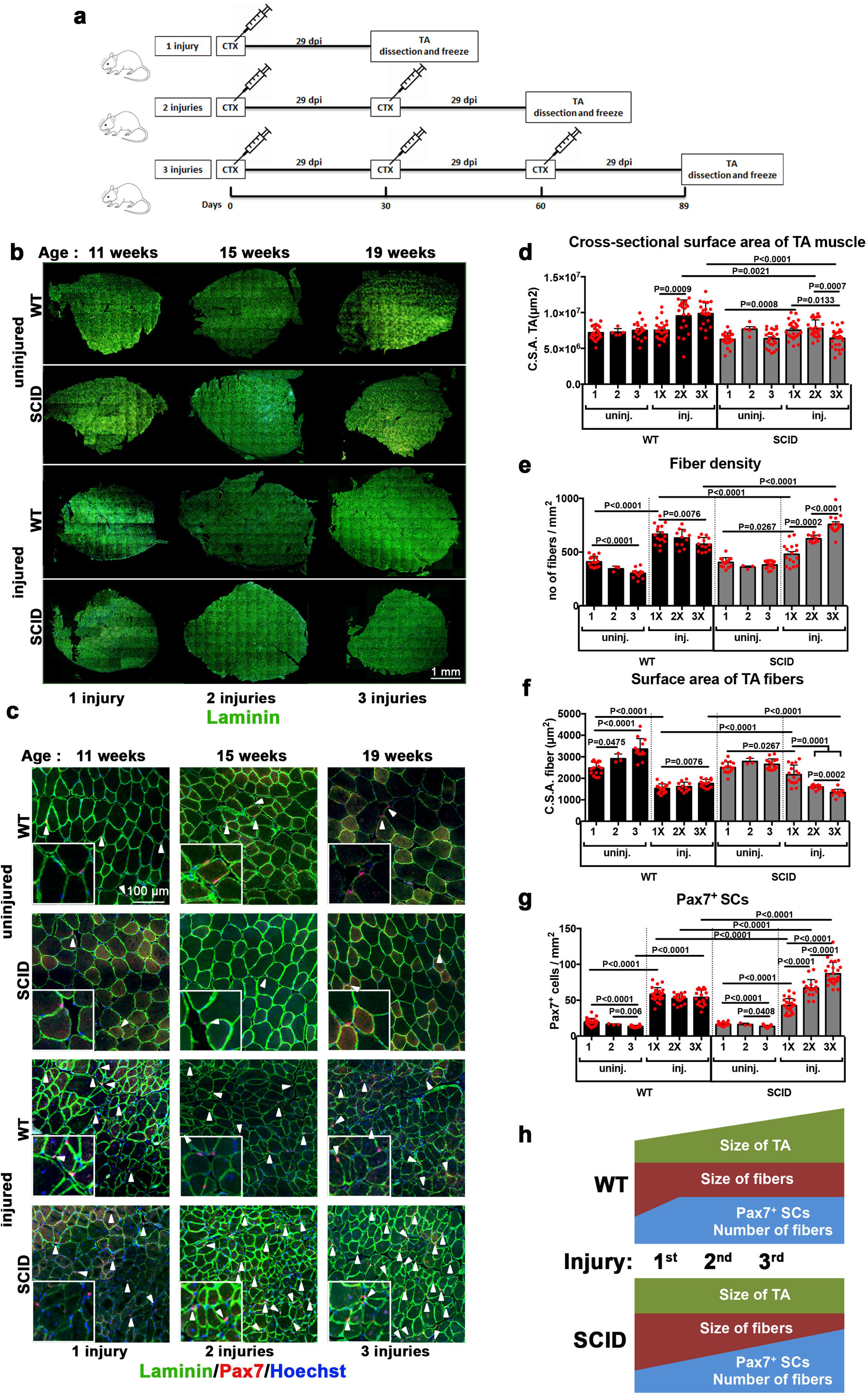
Marked decrease of muscle mass and increased SC population in SCID mice after multiple regeneration events. **a)** Experimental planning of repetitive cycles of muscle injury and regeneration. WT and SCID mice were injected 1 or 2 or 3 times with cardiotoxin, at intervals of 29 days between injections. TA muscles were dissected and frozen either at 29 days post-injury (1 injury), or 59 days post-injury (2 injuries), or 89 days post injury (3 injuries). **b)** Complete sections and **c**) zoomed images of cross sectional TA, immunostained for Laminin (green) and Pax7 (red), and counterstained for Hoechst (blue, nuclei). Histograms of **d)** cross sectional surface area of the entire TA (measured from sections as in Fig. 5d; n=4-5 TA (mice)/condition), **e)** number of fibers per mm^2^ (*i.e.* fiber density) **f)** cross sectional surface area of fibers **g)** number of Pax7^+^ SCs per mm^2^. n=4-5 mice/condition, 5 sections/condition, mean ± SD. Significant P values between conditions according to the Mann Whitney test are indicated on the histograms. **h)** schematic representation (not at scale) of changes in the parameters measured in panels d-g upon each injury/regeneration event per mm^2^ of a TA muscle.

The total body weight of uninjured WT and SCID mice increased from the beginning to the end of the experiment (from 7 to 19 weeks; not shown), but the weight of their TA did not (Table S1). In contrast, after the third regeneration event, TA muscles increased by 34% and 21% in weight in WT and SCID mice, respectively. Overall, the CSA of the TA was smaller in SCID compared to WT mice after the second and third regeneration cycles (Fig. 5b-d). Thus, despite the increasing in body weight, the CSA of the TA muscle was essentially unchanged in SCID mice after 3 regeneration cycles.

The number of fibers (and thereby fiber density), which increased in both mice in the acute phase of the first regeneration (at 5dpi) (see Fig. 4f) remained high in WT mice at the end of the first regeneration cycle, and slightly decreased in the successive ones (Fig. 5e). Conversely, in SCID mice the fiber density increased slowly at the end of the first regeneration cycle, and progressively increased during the successive regeneration cycles. Thus, after the third regeneration event the fiber density was higher in SCID than WT mice.

The fiber size also decreased at 5 dpi (Fig. 4g), and remained low until the third regeneration cycle in all mice, in particular SCID (Fig. 5f). Finally, the density of Pax7^+^ SCs increased from 19/mm^2^ in uninjured mice to 58/mm^2^ after the first regeneration cycle, and remained stable until the third regeneration cycle in WT mice. Conversely, the density of Pax7^+^ SCs kept increasing in SCID mice, and reached 87/mm^2^ after the third regeneration cycle, thus exceeding the number in the WT (Fig. 5g).

In summary, following multiple rounds of injuries, the TA muscle of WT mice was larger in size, and SCID mice of smaller size, compared to uninjured muscle (see Fig. 5h and Fig. 6a). Despite these differences, upon repetitive injuries the TA of either mice contained more fibers, which were of smaller size, and had a higher number of SCs compared to the corresponding uninjured muscle, pointing to different regeneration dynamics in mice mutant for DNAPK.

**Figure 6.**
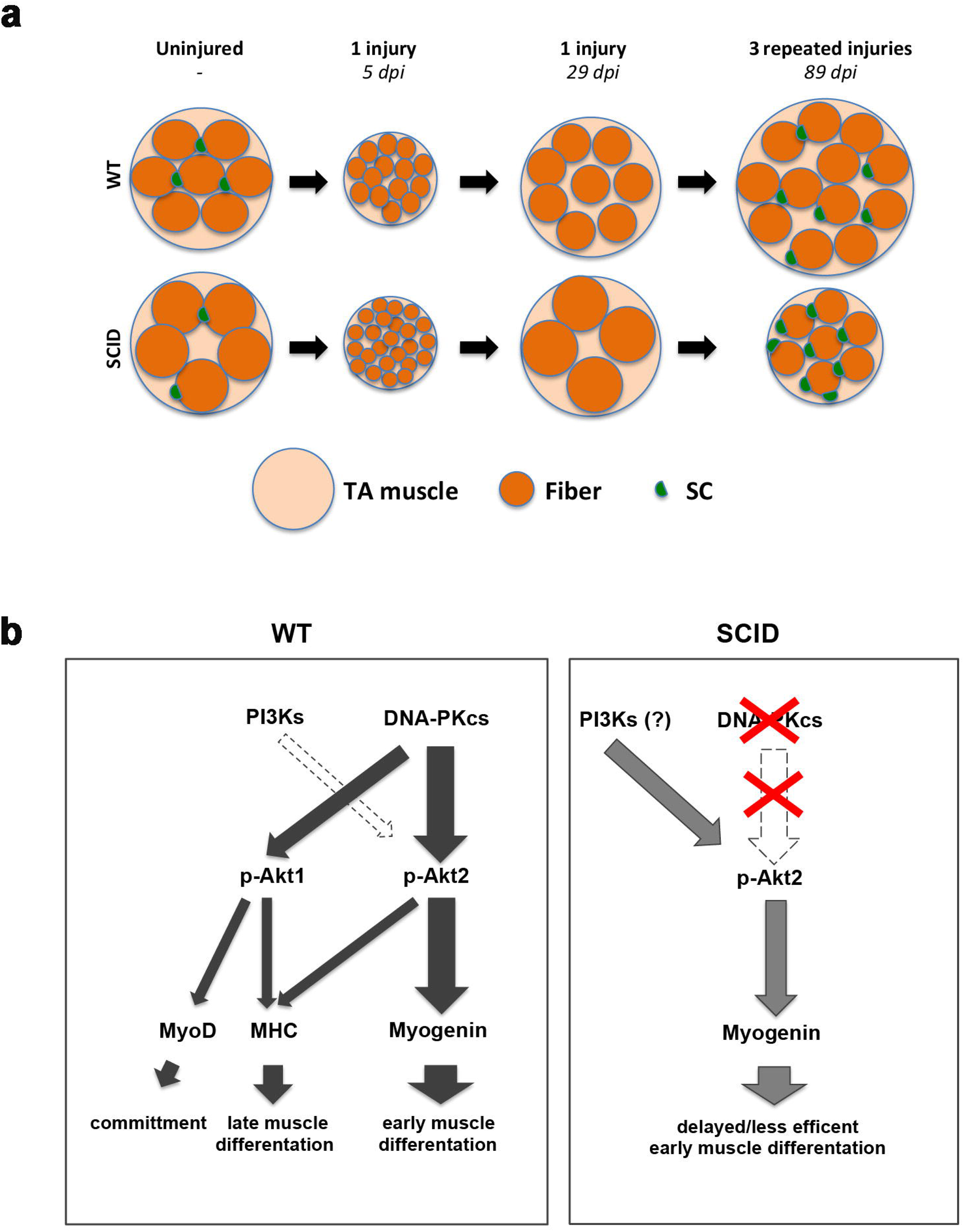
Scheme activation of Myogenin and muscle fibers before and after regeneration in WT vs SCID. **a)** Scheme of cross-sectional area of uninjured (the comparable cross-sectional surface area of uninjured TA muscle as WT: 5.4 ± 1.2mm^2^ and SCID: 5.5 ± 1.1mm^2^), one-time injured TA at 5dpi and 29dpi, and 3 times injured (every 29 days) TA muscle at 89dpi from WT and SCID mice. Big cantaloupe circles represent TA muscle, orange circles represent individual fibers and green half circles represents SCs. The scheme, which is not at scale, aims to summarize the difference in SC number, fiber number, and size in tested conditions. **b)** Left panel: schematic representation of the activation of Myogenin by DNA-PKcs through phosphorylation of Akt2. The activation of other myogenic and differentiation factors (MyoD and MHC) by DNA-PKcs through Akt2 and/or Akt1 is also shown. The larger the arrow the stronger the effect. The minor role of PI3Ks is indicated with a white dotted arrow. In the absence of DNA-PKcs, PI3Ks in large part activates Myogenin (right panel).

## Discussion

Muscle stem cell differentiation undergoes multiple regulatory states. We identify a new regulatory function for DNA-PKcs kinase in this process where it acts independently of its role in DSB repair and *via* activation of Akts. We also show that chronic impairment of this pathway *in vivo* leads to alternative activation of Akts that results in compromised muscle regeneration.

### DNA-PKcs activates the expression of Myogenin in an Akt2-dependent manner

We report here that differentiation of myogenic cells requires the protein kinase DNA-PKcs to promote expression of the key differentiation factor Myogenin. Inhibition or silencing of DNA-PKcs blocks the expression of Myogenin and myogenic differentiation in an Akt2-dependent manner (Fig. 6b), and reduces phosphorylation of Akt. Further, inhibition of Akts as well as Akt2 silencing blocks Myogenin expression. Consistently with these findings, Myogenin expression has been reported to correlate with the expression and activation of Akt2 [30, 31]. Importantly, our findings show that DNA-PKcs acts upstream of Akt2 and is directly responsable for Akt2 activation.

Akts have been shown to play a role in myogenesis upon activation by PI3Ks, specifically Akt1 which promotes SC activation, whereas Akt2 is implicated in differentiation [32], [11], [12, 13]. These studies mostly focused on activated or committed myogenic cells (MyoD^+^). Our findings focus on the pivotal role of regulating the expression of Myogenin and differentiation, which we show requires Akt2 phopshorylation by DNA-PKcs. Conversely, PI3Ks, which are major effectors of Akt activation [33], and were hypothesized to act also during this phase of myogenesis [13], do play a minor role.

Notably, we show that DNA-PKcs intervenes at multiples phases of muscle differentiation, since it acts also upstream and downstream of its major target Myogenin. Indeed, by phosphorylating Akt1, DNA-PKcs promotes expression of MyoD, and by phosphoryating Akt1 and Akt2, it leads to the expression of the late differentiation factor MHC (Fig. 6b). However, inhibition or silencing of DNA-PKcs reduces, but does not block the expression of MyoD and MHC, in contrast to the case for Myogenin.

### DNA-PKcs activates Myogenin expression in the absence of induced DNA damage

DNA-PKcs has been reported to activate Akts essentially at DSB sites where it regulates the DNA repair mechanism of NHEJ [34, 35]. DNA-PKcs is a multifunctional kinase implicated in several processes linked to the repair of DNA damage, in addition to its direct function in NHEJ [36]. Namely, this kinase modulates transcription through interaction with the transcriptional machinery (in particular RNAPII), and phosphorylation of transcription factors including master regulator genes [37], and histones [38]. In these processes, DNA-PKcs is essentially activated by DNA damage [39]. In other cases, for instance, activation of the hypoxia factor HIF1α, the action of DNA-PKcs depends on histone acetylation rather than DNA damage [40, 41].

We identify a novel role of DNA-PKcs in muscle differentiation that we propose to be independent of DNA damage, either induced or of endogenous origin. To activate Myogenin expression, DNA-PKcs does not require irradiation, and this activation is not associated with increased DNA damage or γH2AX levels. Accordingly, the Myogenin promoter does not undergo DNA damage during muscle differentiation [5]. DNA-PKcs-dependent activation of Myogenin appears also to be independent of DNA damage specifically associated with muscle differentiation, and which is induced by a caspase-dependent nuclease (CAD) [5]. In that context caspase-3 and caspase-9 are involved in muscle differentiation [42, 43], and DNA-PKcs phosphorylates the histone H2AX in response to caspase-dependent DNA damage [25]. Transient DNA damage has been proposed to play a role in muscle differentiation, but this process has not been fully elucidated. As such, caspase-induced DNA damage is repaired, likely by nucleotide excision repair (NER), and it is thought to regulate the expression of genes that are relevant for myogenesis, for instance p21 that is necessary for cell cycle withdrawal and thus myogenic differentiation [5, 44]. In our experiments inhibition of caspases did not reduce the number of Myog^+^ cells, whereas inhibition of DNA-PKcs fully depleted this myogenic population. These data show that DNA-PKcs-dependent myogenesis through activation of Myogenin is well distinct from DNA-PKcs phosphorylation of H2AX in response to caspase-induced DNA damage.

### DNA-PKcs interacts with the complex that activates the Myog promoter

DNA-PKcs-dependent expression of Myogenin does not require the presence of the kinase directly on the *Myog* promoter, compatible with the notion that activation is not triggered by DNA damage. Instead, DNA-PKcs interacts with components of the complex that activates *Myog* expression and includes the transcription activator p300 and the transcription factor MyoD [21]. Co-immunoprecipitation experiments showed that this complex also includes Akt1, and Akt2 (at least in complex with p300), in analogy with Akts phosphorylating p300 to promote the interaction with MyoD [21]. Our experiments also indicate that DNA-PKcs itself interacts with Akt1and Akt2. It is thus likely that DNA-PKcs phosphorylates Akt2 that in turn activates p300 and MyoD at the Myogenin promoter. This mechanism appears similar to activation of the fatty acid synthase promoter, although in that case DNA-PKcs activation largely relies on DNA damage at the target promoter [45]. It is possible that DNA-PKcs also activates Akt1 at the MyoD promoter, and Akt1/Akt2 at the MHC promoter in a similar way, although we have not dissected these two mechanisms where DNA-PKcs has a lesser impact.

### Delayed muscle differentiation in SCID mice

Although inhibition and silencing of DNA-PKcs activity severely affects myogenesis in WT SCs in vivo, DNA-PKcs-deficient (SCID) mice do develop muscles. Muscle is normally formed also in enginereed DNA-PKcs knock-out mice that are deleted for the kinase domain [46, 47]. We showed that nevertheless, SCID mice exhibited detectable defects in differentiation *in vivo* and *in vitro* (*e.g.* delayed myotube formation and reduced number of MHC^+^ cells). This was accompanied by a higher number of renewing SCs, as well as MHC^+^ and Myogenin^+^ cells in SCID compared to WT during early muscle regeneration. An increased number of myogenic and differentiated cells in regenerating SCID may result from hyperactivity or, alternatively, delayed regeneration. The latter possibility is in agreement with *in vitro* data reported in this study, which indicate that muscle regeneration is SCID mice is delayed according to the scheme provided by Murphy et al [48].

Notably, regeneration deficits were exacerbated after multiple rounds of muscle injury and regeneration, resulting in smaller TA muscles in SCID mice. SCID TA also contained less myofibers than WT. These alterations suggest that regeneration defects in SCID mice may affect long-term muscle function. Observations that SCID mice regenerate muscle after multiple rounds of injury suggest that DNA-PKcs-deficient mice rely on alternative mechanisms, probably resulting from adaptation to chronic deficiency of DNA-PKcs. These mechanisms would ensure activation of Akt kinases and Myogenin expression. PI3K kinases are good candidates for this task since insulin, that activates PI3Ks, partially rescued the pAkt signal and Myogenin expression in DNA-PKcs-inhibited cells. Moreover, inhibition of PI3Ks to some extent reduced Akt2 phosphorylation and Myogenin levels.

DNA-PKcs deficiency in SCID mice [49] is due to a point mutation (Tyr-4046) that generates an early termination codon, abolishing the kinase activity [50]. The missing domain is thought to be responsible for the stability and/or the folding of the protein, and indeed DNA-PKcs protein levels are negligible in SCID mice [51, 52]. It is unlikely that the residual DNA-PKcs (if any) would trigger myogenesis in SCID mice, since the myogenic function of DNA-PKcs depends on its kinase activity that is missing in these mutants [50]. The kinase domain is also depleted in engineered DNA-PKcs mutants, that also do form muscle [46, 47]. Thus, alternative, PI3K-dependent mechanisms likely promote myogenesis when DNA-PKcs is inactive.

SCID mice are a common model for muscle dystrophy and transplantation studies. For example, SCID/*mdx* mice are immunodeficient due to the DNA-PKcs mutation (NHEJ is required for immunoglobulin V(D)J recombination) and a model of muscular dystrophy due to mutation in the dystrophin gene (*DMD*, Duchenne muscular dystrophy). Immunodeficiency makes these mice good recipients for transplantation experiments, thereafter most studies are concentrated on the effects of immunodeficiency over muscle regeneration, by assessing transplanted cells derived from WT mice or human [53], or focusing on inflammation [54] or fibrosis during regeneration of the injured muscle [55]. Although SCID mice remain a good model for many studies, we show here that impairment of DNA-PKcs results in different muscle regeneration dynamics and muscle composition compared to WT, and these differences should be considered in future studies with these mice.

## Conclusions

We have demonstrated that DNA-PKcs is essential for myogenesis *in vitro* where it activates Myogenin expression through phosporylation of Akt2. To a minor extent DNA-PKcs also activates the upstream myogenic factor MyoD and the downstream marker MHC, thereby playing multiple roles during muscle commitment and differentiation. In the absence of DNA-PKcs, Myogenin is activated by PI3Ks, which however results in delayed dynamics of regeneration. These alterations are exacerbated following repeated rounds of muscle injury. SCID mice that are naturally mutated in DNA-PKcs may have developed compensatory mechanism that may not necessarely operate in WT mice. Since SCID mice are currently used for stem cell transplantation studies, it should be considered that these mice regenerate muscle through an adaptative and essentially DNA-PKcs-independent mechanism.

## Acknowledgements

This work was supported by AFM (research grant (16580), and thesis grant (18425)). We thank the lab of Shahragim Tajbakhsh for the material provided to perform *in vivo* muscle injuries and antibodies targeting myogenic factors, Hiroshi Sakai for guidance to initate the *in vivo* injury experiments, and Shahragim Tajbakhsh for helpful discussion, Sebastien Mella for his advice on statistical analysis. We thank the Virus and Immunity Laboratory at Institut Pasteur (director Dr. Schwartz) for quantification of p24 viral particles, and the center for Translational Science (CRT)-Cytometry and Biomarkers Unit and Phtonic BioImaging Unit of Technology and Service (CBUTechS and PBI UTechS) at Institut Pasteur for technical support in this study.

## Materials & Methods

### Animal Models

For most *in-vitro* experiments and drug treatments SCs were isolated from the limbs of *Tg:Pax7-nGFP* mice [56]. DNAPK-deficient SCs and TA cryosections were obtained from CB17/lcr-Prkdc^scid^/lcrlcoCrl (SCID, severe combined immunodeficiency) mice and Balb/cAnN (wild-type, WT, control) purchased from Charles River laboratories. For regeneration analysis of injured TA muscles, also Rag2^-/-^ γc^-/-^ and C57BL/B6 mice as control were used, provided by Shahragim Tajbakhsh, Paris. The genotype, phenotype and genetic background of each mouse strain are indicated in Table S2.

### Ethics statement

Animals housing, husbandry and handling were performed in the animal facility of Institut Pasteur in accordance with the national and European community guidelines. SC isolations were performed from healthy 7-10 weeks old male mice, *in vivo* injury experiments were performed on healthy 7-19 weeks old male mice according to national and European guidelines, and protocols were approved by the ethics committee at Institut Pasteur and the French Ministry.

### Satellite cell isolation by FACS and cell culture

SCs were freshly isolated from fore and hind limbs of *ad hoc* mice that express the Pax7 marker linked to the fluorescent label GFP (*Tg:Pax7-nGFP* mice), following the isolation protocol, as previoulsy described [57], and in more detail in [58]. Briefly, initially the limbs were chopped off, and precipitated in DMEM Glutamax (GIBCO) with 1% Penicillin Streptomycin (Pen/Strep). The floating lipid particles were successively removed by discarding the excess medium. This step was followed by digestion step with 0.1% Collagenase D (Roche), 0.25% Trpysin (GIBCO) and 0.1 mg/ml DNase I (Roche) in DMEM Glutamax (containing 1% Pen/Strep) for 30 min at 37°C, which was repeated 5 times until full digestion of the tissues. After each digestion step, where the tissue was placed in a foetal bovine serum (FBS) solution to inactive trypsin, the digested material (suspension) was filtered on ice through subsequently 100 µm, 70 µm, and 40 µm cell strainers respectively after each digestion step into foetal bovine serum (FBS) to inactive trypsin. The filtered digests were then pre-spinned for 10 min at 50 x g and the supernatant was collected. The supernatant was then centrifuged for 15 minutes at 550 x g at 4°C. The supernatant was discarded and the pellet resuspended in fresh DMEM Glutamax with 1% of Pen/Strep. Centrifugation was repeated 3 to 4 times until the resuspension was sufficiently transparent to proceed further. After the final spin, the pellet was resuspended in 600 µl of DMEM Glutamax (with 1% Pen/Strep) and 2% FBS. Finally, the samples were processed by FACS (ARIA III, BD Biosciences) to sort SCs, which are Pax7-nGFP^+^, using GFP as marker.

In the case of SCID mice and the corresponding Balb/cAnN control mice, as well as Rag2^-/-^ γc^-/-^ and the control C57BL/B6 mice, which SCs are not *Pax7-nGFP^+^*, the muscle bulk was digested with 2.4 U/ml of Dispase II (Roche), 0.2% of Collagenase A (Roche) and 0.1 mg/ml DNase I (Roche) in DMEM Glutamax (+1% Pen/Strep) for 2 hours at 37°C. Then filtration and centrifugation were performed as indicated above. At the final step, cells were labelled with receptor markers (conjugated antibodies) for 30 min at 4°C. Cells were then sorted with FACS ARIA III, gating the SC population for Sca1^-^, CD45^-^ and CD31^-^, CD34^+^, and Itgα-7^+^, as indicated in the published protocol [58].

### Cell culture

Immediately after sorting, collected SCs were seeded at a density of 5000 cells/cm^2^ in Lab-Tek^TM^ chamber slides (Nunc) or cell dishes pre-coated with 0.1 mg/ml of poly-D Lysine (SIGMA) and Matrigel (Corning). The medium used consisted of: 40% of DMEM Glutamax with 1% Pen/Strep, 40% OF MCDB (SIGMA), 20% of FBS, 2% of Ultraser G (Pall), and 10 µg/ml of insulin when indicated. Cells were incubated at 37°C with 5% CO_2_ and 20% O_2_.

C2C7 cells (immortalised myoblast cell line) [59, 60] were incubated in the same condition as SCs without pre-coating in DMEM Glutamax with 1% Pen/Strep and 20% FBS.

### Inhibitor treatment and irradiation

Cells were pre-treated by addition of inhibitors to the culture medium for 1 hour before irradiation, at a final concentration of 10 µM of the DNAPK inhibitor (= DNAPKi: NU7441) (Axon Medchem), 5 µM of the Akt inhibitor (AKTi, MK2206) (APExBIO), 10 µM of the PI3K inhibitor (LY294002), or 1 µM of the PI3K inhibitor (ZSTK474) (Selleckchem), 20µM of caspase inhibitor Q-VD-OPh (A1901) (APExBIO) or the corresponding volume of vehicle (DMSO, diluent of the drugs). After 1h treatment, cells were irradiated at 5 Gy (with the Xstrahl RS320 Irradiator for X-ray), which is expected to induce approximately 200 DSBs [61] or were not exposed to irradiation (control and vehicle). Cells were incubated for different periods (from 1h to 7 days) post-irradiation before assessing SCs/C2C7 cells differentiation, viability, and cell number. Two experimental plans were used for C2C7 cells (see figures 1, 2, S2, S3). In the first plan, cells were treated with inhibitor (or vehicle) 12h after seeding then analysed at 1-5 or 7 days post seeding (dps). In the second plan, cells were treated with inhibitor (or vehicle) 2.5 or 4.5 days after seeding then analysed at 5 or 7 dps. In experiment in fig.1, cells were treated after overnight seeding and analysed at the indicated times; in the following experiments SCs were treated 5.5 dps and analysed at 7.5 dps.

### shRNA cloning, transfection, and transduction

Lentiviruses were produced from the shRNA plasmids or scrambled (SCR) shRNA for control (Mission shRNA library, Sigma) cloned in *Escherichia coli* (Table S3). ShRNA plasmids, that carry ampicillin and puromycin resistance genes, were amplified in *E. coli* grown overnight in Luria Bertani (LB) broth medium with 100 µg/ml Ampicillin. ShRNA plasmids were extracted and concentrated with NulceoBond® Xtra Maxi Plus kit, Plasmid DNA purification kit (Macherey-Nagel, ref.: 740416.10) following the manufacturer’s protocol. Purified shRNA plasmids and plasmid coding for the viral particle (psPAX2-packaging, pMD2.G-envelope) were co-transfected in HEK-293T cells treated with 25 µM of chloroquine diphosphate (Sigma), a DNA intercalator that increases the transfection efficiency, using the calcium phosphate transfection method [62]. Two days post transfection, the HEK-293T cell culture medium was harvested, filtered through 0.45µm filters, and concentrated by ultra-centrifugation at 19,000 rpm for 90 min at 4°C. After centrifugation the supernatant was discarded and the pellet resuspended in phosphate buffered saline (PBS) in a final volume of 250 µl. For titration purposes, p24 viral particles were quantified by the Virus and Immunity laboratory at Institut Pasteur.

Freshly FACS-sorted SCs were put in culture for 12h then re-seeded at a density of 5,000 cells/cm^2^, and 12h later transduced with shRNAs or SCR (scrambled non-targeting shRNA) concentrated viral particles. The viral particles were added to the cell culture medium with 8 µg/ml of the transduction enhancer Polybrene (hexadimethrine bromide) (Sigma). Six to 8 hours post transduction the culture medium was refreshed (without viral particles and Polybrene), and 48 hours post transduction (or 2.5 dps, considering re-seeding as the starting point), was again refreshed with 1 µg/ml puromycin containing medium for selection of cells stably expressing the shRNA. After 3 days of puromycin selection, *i.e.* 5 days post transduction (corresponding to 5.5 dps), surviving cells were harvested. In parallel, a first sample of cells was harvested 2.5dps and a second sample 5.5dps, in the absence of puromycin treatment and shRNA transduction (control 1 and control 2, respectively).

### Muscle injury

For regeneration analysis, mice were anesthesised with 0.5% Imalgene + 2% Rompun and tibialis anterior (TA) muscles were injected with 50 µl of 10 µM protein kinase C specific inhibitor, cardiotoxin (Latoxan) as previously described [63]. The TA muscles were harvested for analysis 5 days post injury. For repetitive injury, the mice were injected with cardiotoxin for 3 times during 89 days (the second and third injections were done 30 and 60 days after the first injection, respectively). For these experiments, TA muscles were dissected and frozen 29 days post-injury (1 injury), or 59 days post-injury (2 injuries), or 89 days post-injury (3 injuries), and analysed all together. The first injury was performed on 7 weeks old mice and TA muscles injured 3 times were dissected from 19 weeks old mice.

### Immunostaining and histology

Satellite cell-derived cultures were fixed in 4% PFA for 5 min and permeabilized with 0.5 % Triton X-100 for 5 min as previously described [57]. Upon permeabilization, samples were treated for antigen blocking with 2 % BSA and 5% serum in PBS for 1h at room temperature. Primary antibodies were incubated overnight at 4 °C, and secondary antibodies (conjugated with fluorophores), and 5 µg/ml Hoechst 33342 (Sigma, ref #14533) (nuclear counterstaining) for 2h at room temperature.

Freshly dissected TA muscles were frozen in liquid nitrogen cooled isopentane for 30 seconds. The frozen TA muscles were cryo-sectioned into 10 µm slices. These sections were fixed with 4% PFA for 15 minutes at room temperature and antigen unmasking was performed by incubating sections in boiling 10 mM citrate buffer pH 6 for 20 mins prior to immunostaining, as described [56]. For regeneration analysis with embryonic Myosin heavy chain (eMHC) specifically, the TA sections were left unfixed and treated with acetone for 10 min at −20 °C prior to immunostaining. All sections were permeabilized with 0.5% Triton X-100 for 5 min and antigen blocking performed with 10% goat serum (inactivated) or 2% goat serum for eMHC staining in PBS and 0.5% Triton X-100 containing PBS for 1h at room temperature. Incubation with primary and secondary antibodies was performed as for cell immunostaining, see above.

### Antibodies

The following antibodies have been used in this study: mouse monoclonal α-Pax7 (AB_528428), mouse monoclonal α−Myogenin (F5D, AB_2146602), mouse monoclonal α-Myosin Heavy chain, MHC (MF20, AB_2147781) and mouse monoclonal α-embryonic MyHC (F1.652, AB_528358) antibodies from Developmental Studies Hybridoma Bank; rabit polyclonal α-Myogenin (sc-576), rabbit polyclonal α-MyoD (sc-304), Rabbit polyclonal α-GAPDH (sc-25778) from Santa Cruz; mouse monoclonal α-Akt2 (5239), rabbit polyclonal α-phospho-Akt2 (Ser474) (8599), rabbit polyclonal α-Akt1 (C73H10), rabbit polyclonal α-phospho-Akt1 (Ser473) (9018), rabbit polyclonal α-phospho-Akt (9271), rabbit polyclonal α-H2AX (2595) from Cell Signaling technology; mouse monoclonal α-phospho-H2AX(Ser139) (05-636) from Millipore; rabbit polyclonal α-53BP1 (NB100-304) from Novusbio; chicken polyclonal-α-GFP (ab13970), rabbit polyclonal-α-DNAPKcs (ab70250) rabbit monoclonal α-Cyclin A2 (ab181591) from abcam; Rabbit α-Laminin (L9393), Propodium Iodide (P4864) from Sigma; goat α-rabbit fluor 555 (A21428), goat α-rabbit fluor 488 (A11034), goat α-mouse fluor 488 (A11029), goat α-mouse fluor 555 (A21425), goat α-mouse fluor cy5 (A10524), goat α-chicken fluor 488 (A11039), goat α-mouse-HRP (31430), goat α-rabbit-HRP (31460) from Invitrogen; hFAB Rhodamine Anti-GAPDH IgG (12004168), Goat α-mouse Starbright blue 700 (12004159) from Biorad; goat α-rabbit CF770 (10078-1) from Biotium; FITC Annexin V Apoptosis Detection kit I (556547) from BD Biosciences.

### Western blotting

Cells were lysed in lysis buffer (5 mM EDTA, 50 mM Tris pH 8, 150 mM NaCl, 0.5 % Triton X-100, 0.1 % SDS) with 2x of cOmplete^TM^ protease inhibitor cocktail (Roche), 1x PhosSTOP (phosphotase inhibitor, Roche) for 30 min on ice. In order to get rid of viscosity due to DNA, the lysed samples were sonicated 5 times for 10 sec (with 10 sec intervals) at medium frequency in the sonicator bath. Samples were denatured in 1x Laemmli with 1/10 β- mercaptoethanol for 5 min at 95°C and seperated in NuPAGE 4-12% Bis-Tris Protein Gels (Invitrogen). The separated proteins were transferred and bound on nitrocellulose membrane by Trans-Blot Turbo system (BIO-RAD) or classical electro-transfer in 1x NuPAGE transfer buffer (with 20% ethanol) bound on an Amersham Protran nitrocellulose membrane (0.45µm porosity) (GE Healthcare, Life Sciences). The membranes were blocked with 5% nonfat dry milk for detection of non-phosphorylated residues, or 5% BSA for detection of phosphorylated proteins in 1x PBS (phosphate buffered saline) with 0.1% Tween 20 for 1 hour at room temperature. Primary antibodies were incubated in the corresponding blocking buffer (milk or BSA in 1x PBS with 0.1% Tween 20) overnight at 4°C followed by 1h incubation with the secondary antibody (conjugated with either peroxidase [HRP] or fluorophore) at room temperature.

### Imaging and enumeration of immunomarkers

Cells were analysed with a fluorescence microscopy Zeiss Axioplan equipped with an Apotome and Axiovision software or Cell Voyager CV1000, confocal scanner box (Yokogawa). γ-H2AX and 53BP1 foci (DSB markers) were enumerated in at least 30-50 cells/condition, from 3 independent experiments. For differentiation analysis, cells positive for one or more myogenic markers were enumerated per condition, and the percentage was evaluated on the total number of nuclei (Hoechst staining), by assessing 2,000-5,000 nuclei/condition and 5-10 fields/condition. The fusion index was calculated with the number of nuclei/myosin heavy chain positive myotubes, by assessing 2,000-5,000 nuclei/condition and 5-10 fields/condition. Image analyses were performed with the ImageJ 2.0 software.

For the histology analysis, 4-10 complete muscle section per condition were analysed. The complete muscle section images were taken by Cell Voyager CV1000, confocal scanner box (Yokogawa) and SP8 resonant scanner (Leica) as 5 stacks and maximum projection were analysed. Image analyses were performed by ImageJ 2.0 software. The actual numbers of eMHC^-^ and eMHC^+^ fibers, Pax7^+^ and Myogenin^+^ nuclei per section were counted. Upon measurement of cross-sectional surface area (C.S.A.) of the TA muscle sections, average cross-sectional surface areas of fibers for each section were calculated by (C.S.A. of TA muscle section/total number of fibers).

Western blots were either exposed on films and scanned, followed analysis of protein signal intensity by ImageJ 2.0 software when needed or expose by Chemidoc (BIORAD) and in this case protein signals were analysed by Imagelab software. The ratio of proteins were calculated by dividing the the signal intensity of the proteins analysed by Image J and protein intensities were normalised by dividing the protein of interest by the intensity of GAPDH of the same condition.

### RNA extraction and RT-qPCR

Total RNA was isolated from cells using the RNAeasy extraction kits (Qiagen) and reverse transcribed with Superscript IV reverse transcriptase (Invitrogen ref #18090010), following the manufactures’s instructions. The reaction mix consisted of 2.5 µM of Oligo d(T)_20_ primer, 0.5mM dNTPs, up to 1µg template RNA, 1x SSIV buffer, 5mM of DTT and 10U/µl of Superscript^TM^ IV Reverse Transcriptase. The cDNAs were quantified by real-time RT-qPCR using the Power SYBR Green Master mix (Applied Biosystems) on a StepOne Plus RealTime PCR system (Applied Biosystems). Myogenin transcripts were amplified using the forward (5’GTGAATGCAACTCCCACAGC) and reverse (5’CGCGAGCAAATGATCTCCTG) primers, and Tbp transcripts using the forward (5’ATCCCAAGCGATTTGCTG) and reverse (5’CCTGTGCACACCATTTTTCC) primers [64].

### Cell counting to assess proliferation

Cells in culture were washed with PBS, trypsinised with 0.05% Trypsin in EDTA for 5 minutes at 37^°^C. Cell counting was performed either 1) with Scepter, Handheld-automated Cell counter (Millipore), which gives the cell count/ml, detected by 60 μm sensors, or 2) manually with the cell counting chambers (KOVA international, ref #87144) and counting under light microscope. When the Scepter was used, at the beginning of each experiment a triple counting was performed to validate the reproducibility of the measurement, then counting was done once/sample. In several cases counting was performed with both methods to validate the cell number.

### CHIP (chromatin immunoprecipitation) and IP (Immunoprecipitation) assays

For ChIP, 10 × 10^6^ C2C7 cells at 2.5 dps were cross-linked in 1% formaldehyde (Sigma-Aldrich, 252549) for 10 min, followed by addition of 125 mM glycine to stop the reaction (5 min). Cells were then washed in PBS, resuspended in lysis buffer (10 mM Tris-HCl (pH 8), 10 mM EDTA, 0.5 mM EGTA, 0.25% (v/v) Triton X-100, and protease and RNase inhibitors) for 5 min on ice. The pellets were then resuspended in 250 mM NaCl, 50 mM Tris-HCl (pH 8), 1 mM EDTA, 0.5 mM EGTA, for 30 min on ice to wash out non-cross-linked material. The resulting pellets were resuspended in 10 mM Tris-HCl (pH 8), 1 mM EDTA, 0.5 mM EGTA, 1% (w/v) SDS and chromatin was sheared by sonication (final average size of 200-500 base pairs, (bp)). The chromatin was diluted to obtain the following buffer composition: 0.1% (w/v) SDS, 1% (v/v) Triton, 0.1% (w/v) sodium deoxycholate, 10 mM Tris-HCl (pH 8), 150 mM NaCl, 1 mM EDTA, 0.5 mM EGTA, and protease and RNase inhibitors.

ChIPs were carried out by incubating 10 to 20 µg of chromatin with 1 µg of either antibody: rabbit polyclonal-α-DNAPKcs (ab70250), rabbit polyclonal α-MyoD (sc-304), or α-rabbit IgG (Santa Cruz Biotechnology, sc-2027) as a control. After overnight incubation at 4 °C, 40 µl of protein G-Dynabeads™ (Thermo-Fisher Scientific, 10004D) were added on lysate for 2 h at 4 °C. After extensive washing, DNA was isolated from the beads by successive boiling for 10 min in the presence of 10% (w/v) Chelex 100 Resin (Bio-Rad, 1421253), incubating at 55 °C for 30 min in the presence of 100 µg.ml^−1^ of Proteinase K (Eurobio Ingen, GEXPRK01-E), and boiling again for 10 min. After centrifugation, the resulting supernatant was used as a direct template for qPCR detection of 5 different regions of Myogenin promoter (Fig. 3c) (primers listed in Table S4). Immunoprecipitated chromatin with the indicated antibodies was calculated as a percentage of the input DNA after normalization with control ChIP performed with rabbit IgGs (Mock) [65].

For the IP assays, 10X10^6^ C2C7 cells were lysed at 2.5dps by lysis buffer for 30 mins on ice as for WB. The 40µl of protein G-Dynabeads™ (Thermo-Fisher Scientific, 10004D) were incubated with primary antibodies (α-mouse IgG (Santa Cruz Biotechnology, sc-2025) and α-rabbit IgG (Santa Cruz Biotechnology, sc-2027) were used as controls) for 2 hrs at 4°C followed by incubation of antibody-conjugated beads with 10µg of cell lysates overnight at 4°C. The immunoprecipitated proteins were analysed by Western blot.

### Statistical analyses

Statistical analyses were performed using the Graphpad Prism (version 6-7) software. For multiple grouped data was applied a multiple comparison analysis: 2-way ANOVA coupled with Dunnett’s multiple comparisons test; for the other experiments 1-way ANOVA with post-hoc Tukey’s multiple comparisons test, non-parametric Mann-Whitney test or Unpaired Student T-test were performed, as indicated.

## Supplementary Tables

**Supplementary Table S1:**
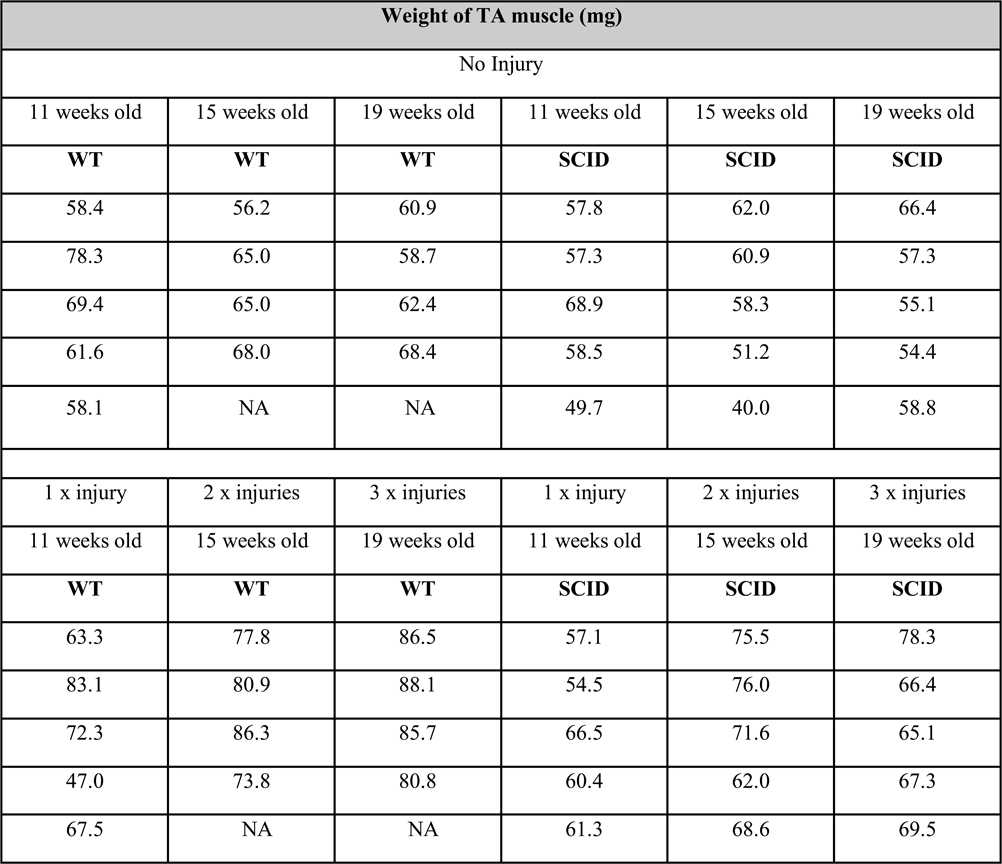
TA muscle weight after 1, 2, or 3 injuries, or uninjured, from WT and SCID mice. NA: not applicable.

**Supplementary Table S2:**
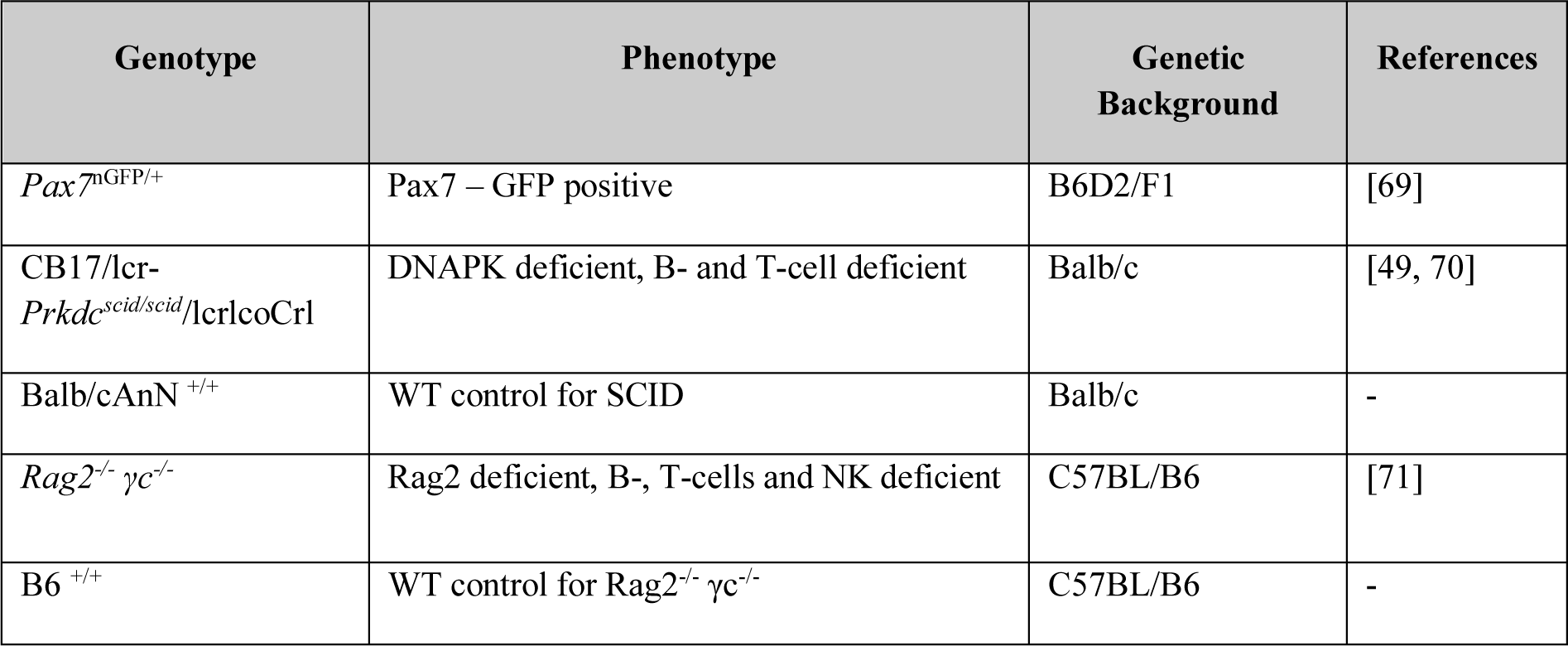
List of mice strains used. The genotype, phenotype, and genetic background of each strain are indicated.

**Supplementary Table S3:** List of shRNAs. Reference numbers, plasmid codes, and target genes are indicated. SCR, scramble.

| Mission shRNA Library (Sigma) | Plasmid | Target gene |
| --- | --- | --- |
| Ref:SHC002 | pLKO.1_Puro control | shRNA (sh Control) (SCR) |
| Ref:TRCN0000023863 | pLKO.1_Puro DNAPKcs | shRNA (sh DNAPKcs-1) |
| Ref:TRCN0000361373 | pLKO.1_Puro DNAPKcs | shRNA (sh DNAPKcs-2) |
| Ref:TRCN0000310882 | pLKO.1_Puro Akt2 | shRNA (sh Akt2-1) |
| Ref:TRCN0000301381 | pLKO.1_Puro Akt2 | shRNA (sh Akt2-2) |
| Ref:TRCN0000304684 | pLKO.1_Puro Akt | shRNA (sh Akt1) |

**Supplementary Table S4:**
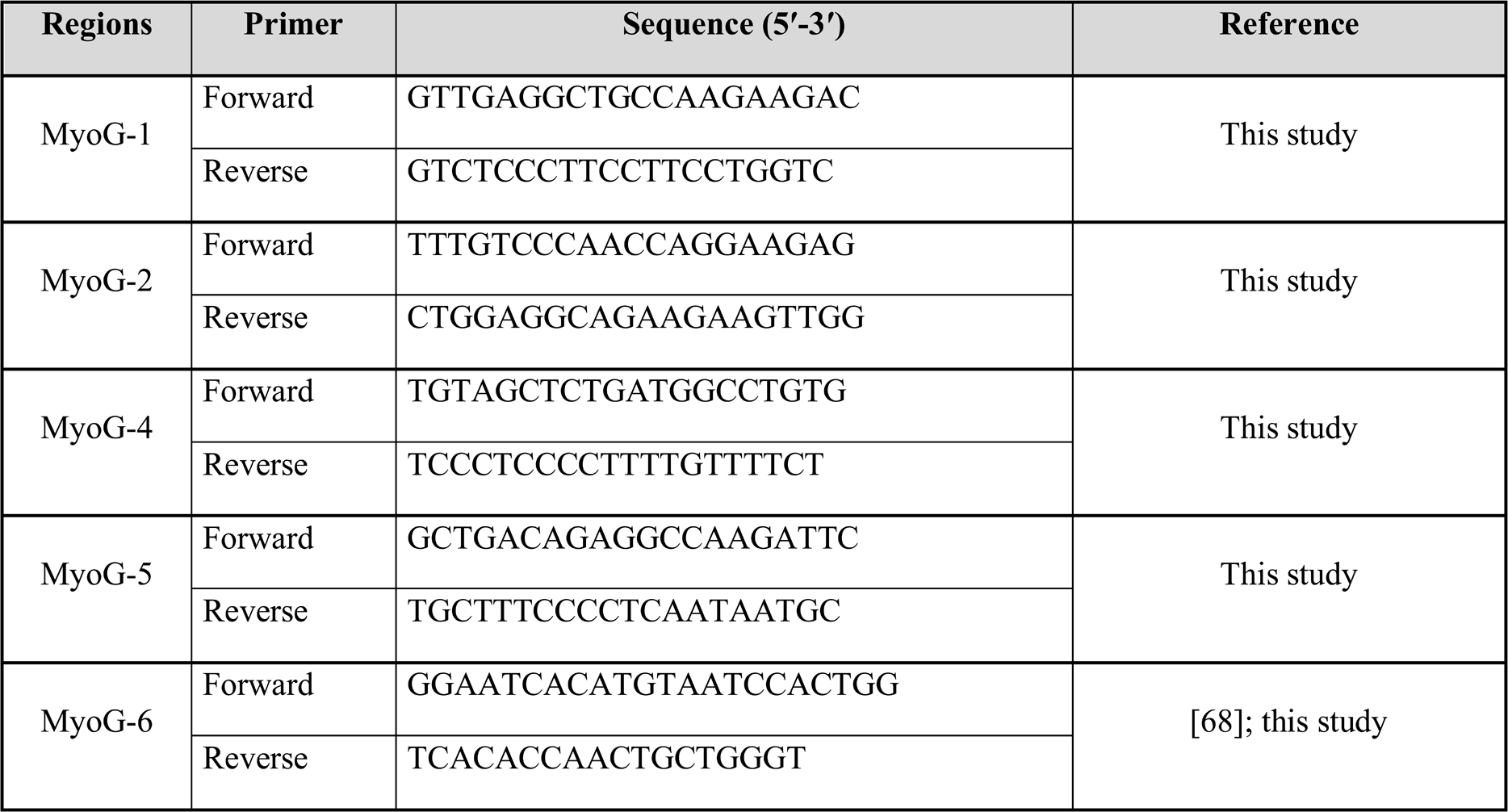
Primer pairs used to assess the promoter region of *MyoG*. The primers have been designed to amplify regions up to 7kbp upstream of the *MyoG* initiation of transcription, where an enhancer region has been previously described [22].

## Supplementary Figure Legends

**Supplementary Figure S1. Inhibition of DNA-PKcs blocks myogenic differentiation of satellite cells in the presence and in the absence of irradiation (induced DSBs)**

**a**) Representative merged images of immunofluorescent labelling of cells 5 days post-IR (and non-irradiated controls): MyoD (red), Myogenin (green), and nuclei counterstained with Hoechst (blue). Representative cells with progressively increased levels of differentiations are indicated with red arrows (MyoD^+^/Myogenin^-^ cells, in red), yellow arrows (MyoD^+^/Myogenin^+^ cells, in yellow), and white arrows (MyoD^-^/Myogenin^+^cells, in green). **b**) Histograms of myogenic differentiation and growth curves of non-irradiated cells ± DNAPKi. Proliferation evaluated with the cell number, and differentiation with immunofluorescence of myogenic markers. Each condition was tested with SC derived from n=3-7 mice, mean ± SD. Myogenic markers have been analysed in 5-10 fields/condition (2,000-5,000 cells), and extrapolated to the total cell number. Significance evaluated by by 2-way ANOVA (F=8.6, DFn=28, DFd=147, p<0.000001), with post-hoc Dunnett’s multiple comparisons test; significant P values are indicated in histogram. P values and bars and are of the same colour as the category that is compared. Panels a-b, experimental conditions as in Fig. 1a. **c**) C2C7 cell number upon irradiation in the presence of increasing doses of DNAPKi, or in the absence of inhibitor. Cells were treated and irradiated at 2.5 dps and harvested at 5 dps. N=3 independent experiments, mean ± SD, Significance by 2-way ANOVA (F=6.20, DFn=4, DFd=22, p<0.0017), with post-hoc Dunnett’s multiple comparisons test against vehicle (DMSO), significant P values are indicated on histogram. The concentration of DNAPKi corresponding to the lowest dose that blocks cell proliferation upon irradiation (10 µM) was used in this study, unless otherwise specified. **d**) Representative images of panel c. **e)** Representative images of SCs cells 4h and 6h post-IR, and non-irradiated controls, immunostained for the DSB repair markers γH2AX (green) and 53BP1 (red), nuclei counterstained with Hoechst (blue), and merge. Efficiency of DSB repair is evaluated by disappearance of repair markers. Histogram report the number of **f)** γH2AX foci/cell (nucleus) and **g)** 53BP1 foci/cell (nucleus) from 4h to 24h post irradiation. Mean ± SEM for each condition. Significance by 1-way ANOVA (γH2AX: F=174, DFn=17, DFd=1892, p<0.0001. 53BP1: F=103.2, DFn=17, DFd=1892, p<0.0001) significant P values by post-hoc Tukey’s multiple comparisons test are indicated on histogram n=30-40 cells/condition. N= 3 independent experiments.

**Supplementary Figure S2. DNA-PKcs inhibitor blocks differentiation of C2C7 cells in the presence and in the absence of irradiation**

**a)** Schematic representation of the experiment. Immortalized myogenic C2C7 cells were pre-treated with DNA-PKi before irradiation (or no-IR) and kept in culture for 1 to 5 days before analysis. **b)** Representative merge images of immunofluorescent labelling of cells 5 days post-IR (and non-irradiated controls). Proliferation was evaluated by the cell number and differentiation by myogenic markers, as in Fig 1. MyoD (red), Myogenin (green), and nuclei are counterstained with Hoechst (blue). Representative cells with progressively increased stage of differentiations are indicated with red arrows (MyoD^+^/Myogenin^-^ cells, in red), yellow arrows (MyoD^+^/Myogenin^+^ cells, in yellow), and white arrows (MyoD^-^/Myogenin^+^cells, in green). Undifferentiated C2C7 myoblasts are originally MyoD^+^. Histograms of myogenic differentiation and growth curves of **c)** non-irradiated and **c)** irradiated C2C7 cells ± DNAPKi. Significance by 2-way ANOVA (Non-irradiated: F=112.8, DFn=11, DFd=47, p<0.0001. Irradiated: F=87.06, DFn=11, DFd=48, p<0.0001), with post-hoc Dunnett’s multiple comparisons test, significant P values are indicated on the histogram. The P values and error bars are of the same colour as the category that is compared. n=3 experiments per condition, mean ± SD. Myogenic markers have been analysed in 5-10 fields/condition (2,000-5,000 cells), and extrapolated to the total cell number.

**Supplementary Figure S3. DNA-PKcs inhibitor affects myogenesis by reducing proliferation and blocking differentiation of myogenic cells**

**a)** Immunolabelling of SC-derived cells at 5.5 dps without DNAPKi treatment (control 1; at this time point cells were essentially myoblasts [88% Myog^-^ and Myosin Heavy Chain, MHC^-^, a structural marker for myotube formation]) and 7.5 dps (with and without [control 2 and vehicle] DNAPKi treatment). Merge of MHC (yellow), Myogenin (red), Pax7-nGFP (green) and Hoechst counterstaining (nuclei, blue). Representative MHC^+^ structures are indicated with a white arrow, Myogenin^+^ cells (red) with a red arrow, and Pax7-GFP^+^ cells (green) with a green arrow. **b)** Histograms display the cell number and differentiation stage of irradiated SC-derived cells treated or not with DNAPKi, and defined by the indicated combinations of myogenic markers (as in Fig. 1d). Myogenic markers have been analysed in 5-10 fields/condition and extrapolated to the total cell number. mean ± SD for each category. Significance by 2-way ANOVA (F=2.53, DFn=9, DFd=32, p=0.0255), with post-hoc Dunnett’s multiple comparisons test; significant P values are indicated in histogram. P values and error bars are of the same colour as the category that is compared. **c)** Fusion indexes of irradiated SC-derived cells by enumeration of nuclei/myotube (as in Fig. 1e). Significance by 2-way ANOVA (F=31.70, DFn=15, DFd=40, p<0.0001), with post-hoc Dunnett’s multiple comparisons test; significant P values are indicated in histogram. P values and error bars are of the same colour as the category that is compared. Significance bars are of the same colour as the category that is compared. n=3 experiments, n=2,500-5,000 cells analysed/condition, mean ± SD. Panels a-c, experimental conditions as in Fig. 1c. Treatment with DNAPKi in committed C2C7 cells (**d-g**) and during differentiation (**h-k**). For simplification, differentiation was evaluated considering Myogenin^+^ *versus* Myogenin^-^ cells. **d**) Schematic representation of the experiment. At 2.5 dps, during mid myogenesis cells were pretreated for 1h with 10 µM of DNAPKi or the corresponding volume of vehicle before irradiation (or were not irradiated), and then analysed at 5 and 7 dps. This experiment mimics the condition of SCs in Fig. 1c, as indicated by comparable ratio of Myog^+^/Myog^-^ cells in control samples (see panel a). Proliferation evaluated with the cell number, and differentiation assessed with immunofluorescence of myogenic markers (MyoD, Myogenin). **e)** Representative merge images of immunofluorescent labelling of cells at 2.5dps and 7dps: MyoD (red), Myogenin (green), nuclei counterstained with Hoechst (blue). Representative cells with progressively increased levels of differentiations are indicated: red arrows (MyoD^+^/Myogenin^-^ cells, in red), yellow arrows (MyoD^+^/Myogenin^+^ cells, in yellow), and white arrows (MyoD^-^/Myogenin^+^cells, in green). Histograms of the cell number and differentiation state of **f)** non-irradiated and **g)** irradiated C2C7 cells treated or not with DNAPKi. The differentiation state is defined by Myogenin immunolabelling: Myogenin^-^ cells [undifferentiated] and Myogenin^+^ [differentiated] cells are shown in the orange and purple part of columns, respectively. Significance by 2-way ANOVA (Non-irradiated: F=7.997, DFn=6, DFd=107, p<0.0001. Irradiated: F=12.68, DFn=6, DFd=103, p<0.0001), with post-hoc Dunnett’s multiple comparisons test, significant P values are indicated in histogram. The P values and error bars are of the same colour as the category that is compared. **h)** Schematic representation of the experiment, which consists in a later stage, with myogenic cells having extensively proliferated, and which are expected to differentiate and form myotubes before treatment. At 4.5 dps, during advanced myogenesis, cells were pretreated for 1h with 10 µM of DNAPKi or the corresponding volume of vehicle before irradiation (or were not irradiated), and then analysed at 7 dps. **i)** Representative merge images of immunofluorescent labelling of cells at 4.5 and 7 dps: the indications are in same colour as in panel e. Histograms of the cell number and differentiation stage of **j)** non-irradiated and **k)** irradiated C2C7 cells treated (or not) with DNAPKi, and defined as in panels c and d. Significance by 2-way ANOVA (Non-irradiated: F=41.38, DFn=3, DFd=98, p<0.0001. Irradiated: F=39.58, DFn=3, DFd=100, p<0.0001), with post-hoc Dunnett’s multiple comparisons test, significant P values are indicated in histogram. The P values and error bars are of the same colour as the category that is compared. n=3 experiments. The Myogenin marker has been analysed analysed in 5-10 fields/condition corresponding to n=2,500-5,000 cells/condition, and extrapolated to the total cell number; mean ± SD.

**Supplementary Figure S4. Regeneration in WT and SCID mice**

**a)** Representative images of un-injured and regenerating TA sections of WT or SCID mice 5 and 7 dpi. Extracellular matrices bordered by laminin (green) and regenerating fibers marked with eMHC (red), nuclei counter-stained with Hoechst; (quantifications are shown in Fig. 5e-g). **b)** Quantification of regenerating eMHC^+^ fibers/mm^2^ on TA sections in WT (C57BL/B6) and immunodeficent *Rag2^-/-^γc^-/-^* mice. n=3 mice, 10 sections/condition, mean ± SD. Significant P values between conditions by Mann Whitney test are indicated on the histograms.

**Supplementary Figure S5. Original gels and blots**

The original blots and the corresponding Red Ponceau stainings are shown for each figure panel. The cropped part is indicated with a rectangle ± the sign of scissors.

