## Supplementary figures and images for "DNA-PKcs regulates myogenesis in an AKT-dependent manner independent of induced DNA damage"

### Supplementary Figures S1 to S5

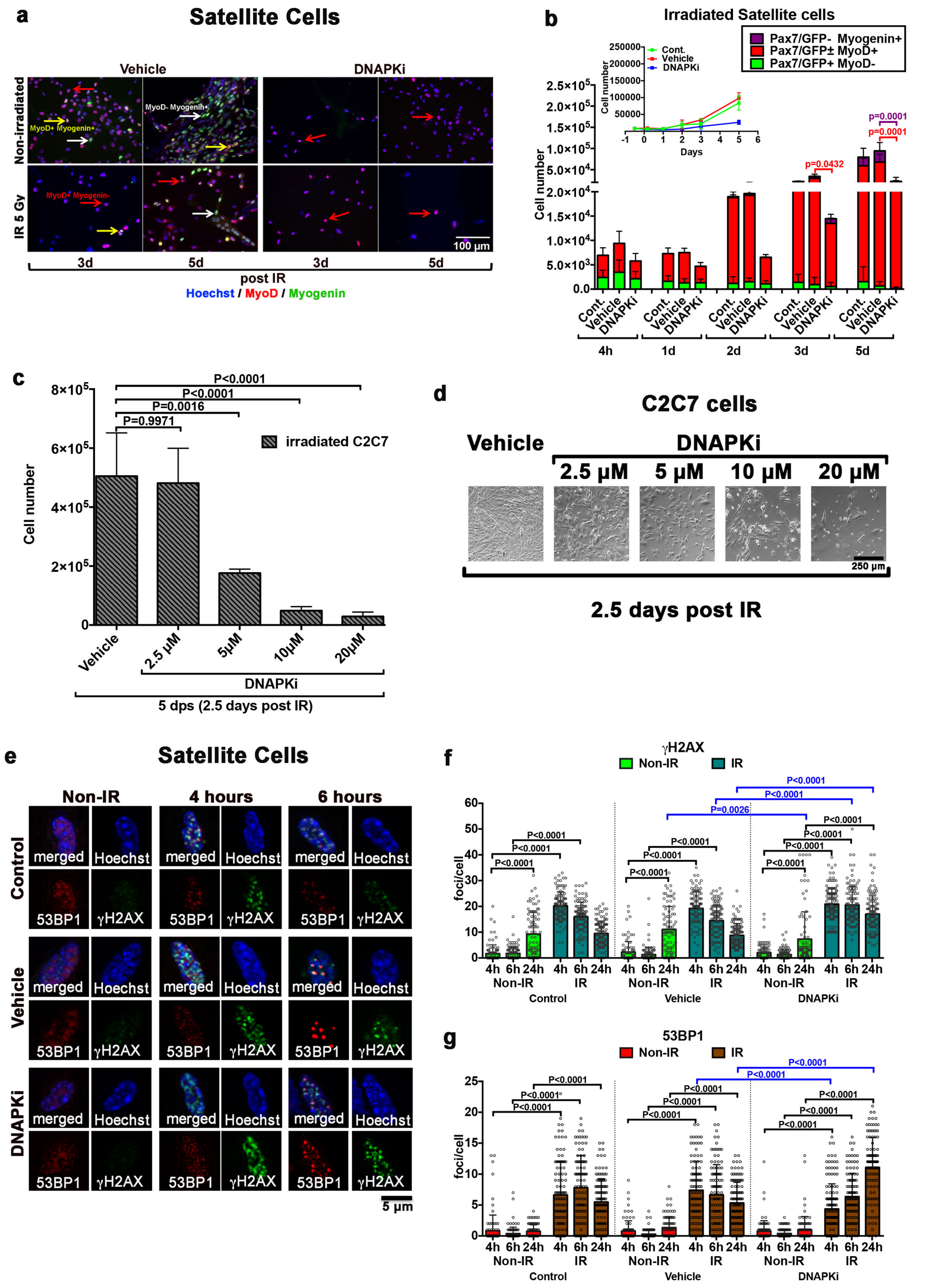

#### C2C7 cells

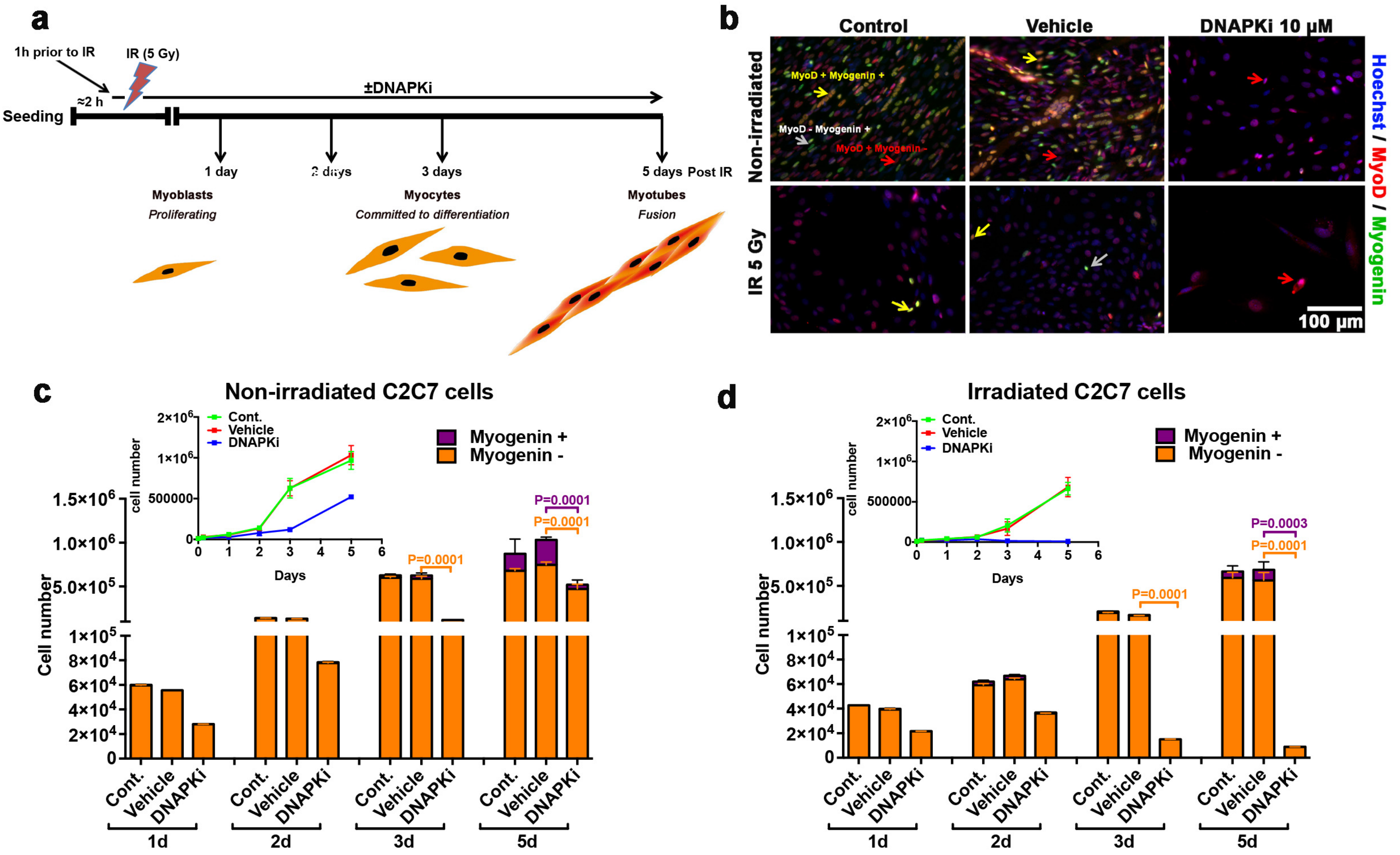


Figure S2

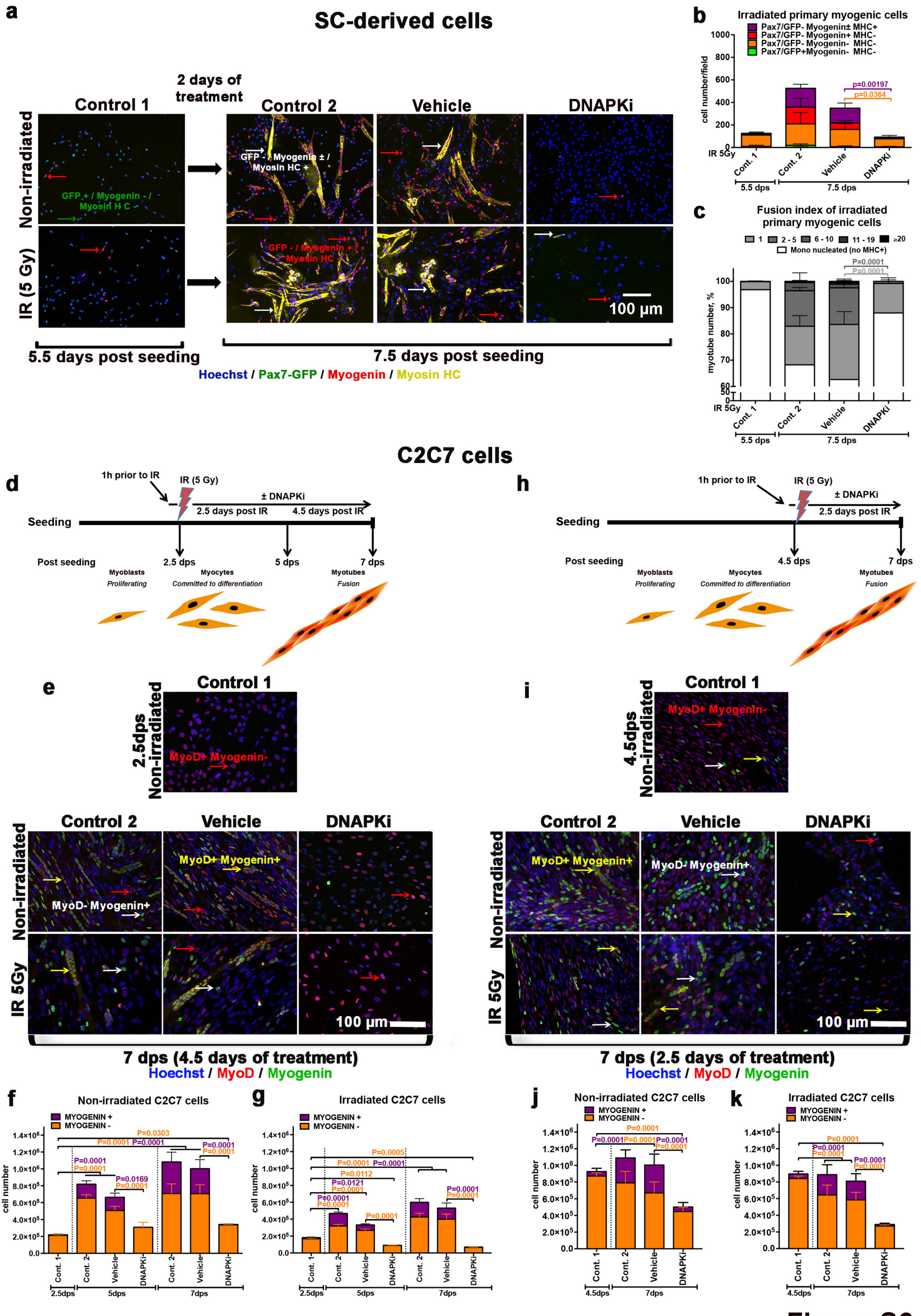

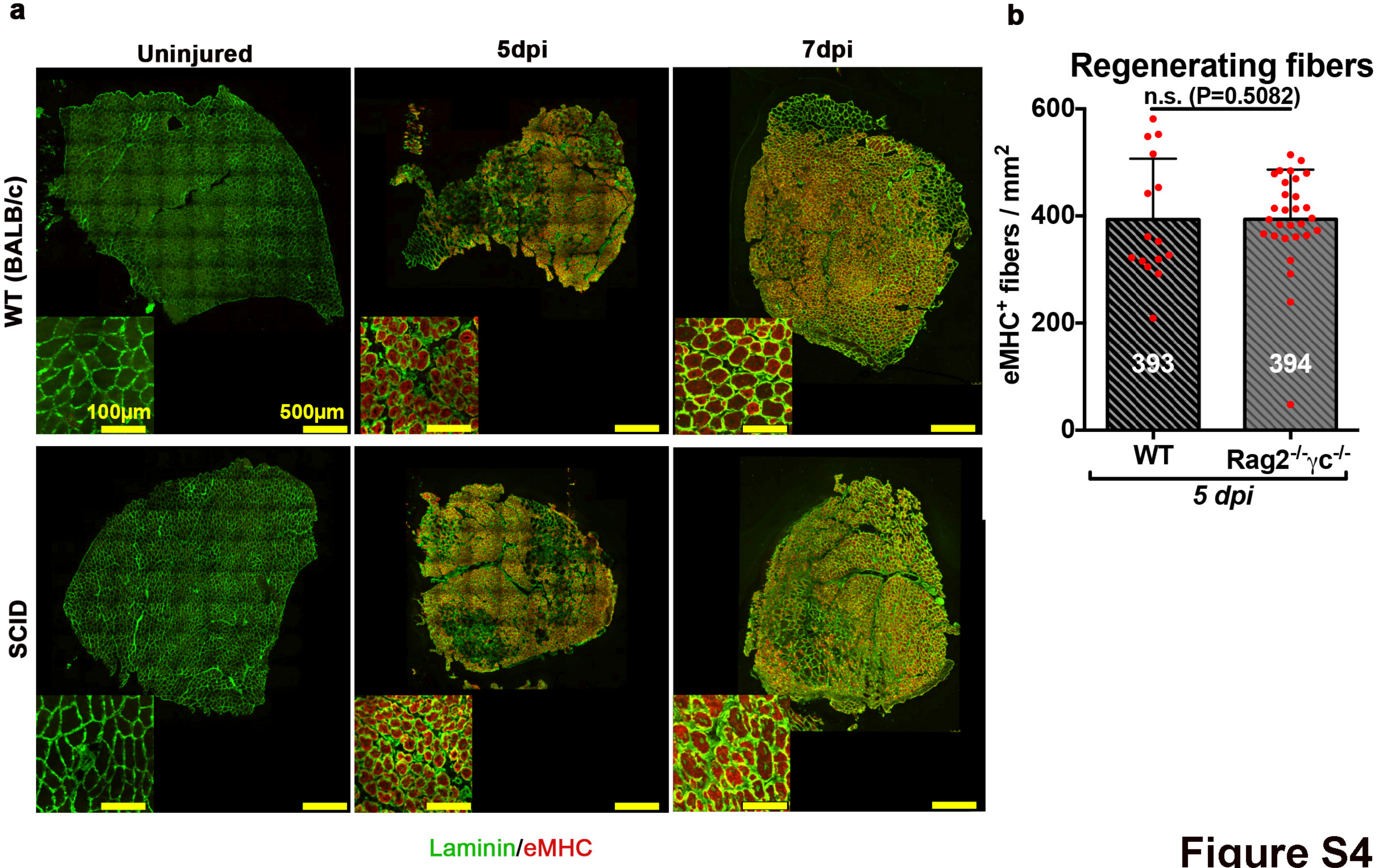

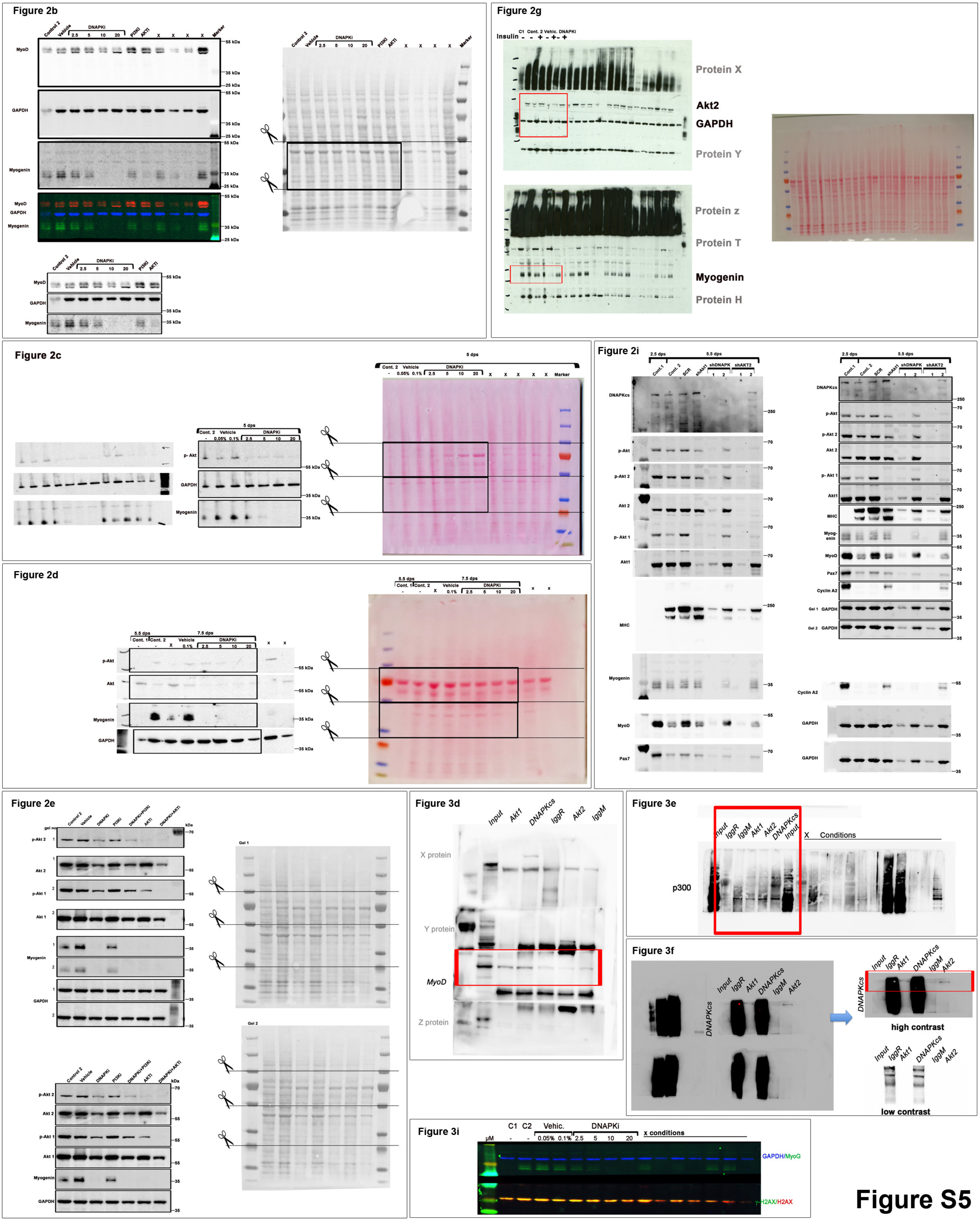
